# Integrative Teaching of Metabolic Modeling and Flux Analysis with Interactive Python Modules

**DOI:** 10.1101/2022.11.16.516820

**Authors:** Joshua A.M. Kaste, Antwan Green, Yair Shachar-Hill

## Abstract

The modeling of rates of biochemical reactions – fluxes – in metabolic networks is widely used for both basic biological research and biotechnological applications. A number of different modeling methods have been developed to estimate and predict fluxes, including kinetic and constraint-based (Metabolic Flux Analysis and Flux Balance Analysis) approaches. Although different resources exist for teaching these methods individually, to-date no resources have been developed to teach these approaches in an integrative way that equips learners with an understanding of each modeling paradigm, how they relate to one another, and the information that can be gleaned from each. We have developed a series of modeling simulations in Python to teach kinetic modeling, Metabolic Control Analysis, l3C-Metabolic Flux Analysis and Flux Balance Analysis. These simulations are presented in a series of interactive notebooks with guided lesson plans and associated lecture notes. Learners assimilate key principles using models of simple metabolic networks by running simulations, generating and using data, and making and validating predictions about the effects of modifying model parameters. We used these simulations as the hands-on computer laboratory component of a four-day metabolic modeling workshop and participant survey results showed improvements in learners’ self-assessed competence and confidence in understanding and applying metabolic modeling techniques after having attended the workshop. The resources provided can be incorporated in their entirety or individually into courses and workshops on bioengineering and metabolic modeling at the undergraduate, graduate, or postgraduate level.

## Introduction

Metabolic modeling provides scientists with a quantitative description of the *in vivo* rates of biochemical reactions in biological networks. These rates of biochemical reactions – fluxes – are a function of many layers of cellular regulation (transcriptional, translational, post-translational, etc.) and relate directly to the living system’s functional phenotype. Understanding metabolic flux thus provides important insights into biological systems and underlies efforts to rationally modify their metabolism to suit our biotechnological needs [1].

Fluxes in metabolic pathways and networks cannot be directly measured, necessitating the use of mathematical modeling approaches to estimate or predict them. These approaches can be broadly categorized into kinetic and constraint-based methods. Within both categories, methods exist both for predicting fluxes and for estimating them from experimental data. Kinetic methods involve simulating the dynamically changing fluxes and metabolite concentrations in a metabolic network over time [2], whereas constraint-based methods like Flux Balance Analysis (FBA) [3] and Metabolic Flux Analysis (MFA) [4] estimate steady-state fluxes using linear optimization principles or experimentally-measured isotopic labeling data.

Metabolic modeling, and particularly constraint-based modeling approaches, have been used productively to aid in biotechnological applications. For example, Metabolic Flux Analysis techniques using isotopic labeling informed the engineering of the bacterium *Corynebacterium glutamicum* to produce high concentrations of lysine [5–7]. Flux Balance Analysis has been deployed to improve the microbial production of a number of bioproducts, including threonine [8] and valine [9], and in ambitious reengineering efforts like that described in [10] where FBA and related methods including [11] were used to enable engineering of normally heterotrophic *Escherichia coli* to incorporate CO2 into its biomass using a heterologously expressed Calvin-Benson Cycle. These and an increasing number of other metabolic modeling applications indicate that this is an area that is of great value to learners and practitioners in biology, biochemistry, and chemical engineering.

Related to kinetic metabolic analysis, Metabolic Control Analysis (MCA) provides mathematical tools for understanding how control over flux and internal metabolite concentrations are distributed between the enzymes in a biochemical network [12,13]. Like metabolic flux modeling and mapping the questions addressed by Metabolic Control Analysis have major biotechnological implications. We believe it therefore makes sense to introduce and teach concepts in MCA along with kinetic and constraint-based metabolic modeling techniques.

Although previous studies have described and provided resources for teaching kinetic metabolic modeling [14], FBA [3,15], MFA [16,17], and MCA [18–20], there are not any published and freely available instructional resources for introducing these toolsets to learners in an integrative and interactive fashion. Moreover, although papers and books exist describing how to experimentally approach 13C-MFA [for example 21-24] or the theoretical background behind the technique [for example 25,26], we are not aware of any dedicated and published educationally focused resources for introducing learners to the theoretical background behind label-assisted MFA. We believe introducing learners to all of these major areas of metabolic modeling together allows them to appreciate their interconnections and better evaluate what approach(es) may be useful to their own research and/or engineering goals than if they encounter them in isolation.

To address this gap in the biochemistry education literature, we developed a series of interactive Python-based Jupyter notebooks featuring exercises that give learners hands-on experience with kinetic modeling, FBA, MFA, and MCA. These notebooks were used as the hands-on laboratory exercises for the 2022 iteration of an annual metabolic modeling workshop at Michigan State University. To assess the efficacy of the workshop and the interactive exercises, surveys were distributed to participants – a mix of graduate students and postdoctoral researchers – before, immediately after, and four months after the workshop to measure self-assessed competence and confidence in metabolic modeling techniques and in the application of these techniques to learners’ own research questions. Although the materials are structured with a particular sequence and timeline, the individual notebooks, paired with appropriate lecture material, contain sufficient explanation to be flexibly incorporated into different course or workshop structures.

## Methods

### Exercise Development

All simulation code was written in Python and packaged and presented in Jupyter notebooks [27]. Numpy [28] and SciPy [29] were used to handle data import and export and calculate control coefficients for MCA. Interactive elements were incorporated into the notebooks using the *ipywidgets* package. MFA simulations were run in Python using the package *mfapy* [30] and FBA simulations were run using *cobrapy* [31]. For the FBA exercises, the genome-scale model of *E. coli’s* metabolic network iJO1366 [32] was used along with a smaller “core” model of *E. coli’s* metabolic network [33]. Several example networks from [25] were adopted for demonstration purposes throughout the notebooks.

Time-courses of metabolite concentrations, fluxes, and labeling were generated in kinetic simulations featuring reversible or irreversible first-order and Michaelis-Menten kinetics. Euler’s method was used to generate all concentration, flux, and labeling values. In most of the simulations that feature labeling, all metabolites are treated as having only one labelable position, so the proportion of labeled and unlabeled metabolite is tracked. In the simulations in the notebook for Day 4 (see **Table 1**), both one- and two-carbon molecules are present, so the quantities of unlabeled, half-labeled, and fully-labeled species for each metabolite are calculated and tracked independently to allow for comparison with 13C-MFA flux map results.

**Table 1:** A table describing the contents of the interactive exercises presented in this publication. Key concepts (as outlined in the text)

| Day | Section(s) | Contents | Concept |
| --- | --- | --- | --- |
| **1** | **1.0 – 1.2** | Introduction to the Jupyterlab Interface. |  |
|  | **2.0 – 2.2** | Exploration of a simulation demonstrating first-order kinetics. |  |
|  | **3.0 – 3.1** | Exercise on inferring kinetic parameters from example datasets. | 1 |
|  | **4.0** | Exercise demonstrating the relationship between model architecture and the information contained in each datapoint. | 1 |
|  | **5.0 – 5.1** | Introduction to metabolic steady-state and the utility of labeling data. |  |
| **2** | **1.0** | Introduction to reversible first-order kinetic models |  |
|  | **2.0** | Exercise on inferring model parameters in the presence of reversibility | 1 |
|  | **3.0 – 3.4** | Exploration of metabolic control analysis, including calculation of flux and concentration control coefficients as well as elasticities. | 3 |
|  | **4.0** | Comparison of results gathered in **3.0 – 3.4** “by hand” with results from an automated MCA script. | 1 |
| **3** | **1.0 – 1.2** | Metabolic control analysis with branching networks, negative control coefficients, and modeling a system with an incomplete network description. | 3 |
|  | **2.0 – 2.2** | Kinetic modeling with Michaelis-Menten kinetics. |  |
|  | **3.0** | Fitting a dataset using either first-order or Michaelis-Menten kinetics in the presence or absence of noise. | 1 |
|  | **4.0** | Kinetic modeling with reversible Michaelis-Menten kinetics. |  |
|  | **5.0** | Using MCA to calculate response coefficients. | 3 |
| **4** | **1.0 – 1.2** | A kinetic simulation that incorporates labeling dynamics, for comparison with <sup>13</sup>C-MFA and FBA. |  |
|  | **2.0** | Introduction to FBA modeling. | 2 |
|  | **3.0 – 3.3** | Introduction to FVA and randomized sampling methods in FBA. | 2 |
|  | **4.0 – 4.2** | Introduction to <sup>13</sup>C-MFA and comparison with results from **1.0 – 1.2**. | 2, 1 |
|  | **5.0** | Discussion about incorporating metabolic modeling into one’s own work and/or research. |  |

### Survey Ethics and Analysis

The survey component of this study was deemed exempt by the Michigan State University Office of Research Regulatory Support. Survey respondents were asked to self-assess their confidence in and understanding of kinetic and constraint-based metabolic modeling methods and the application of these methods to their own research goals on a Likert scale [34]. Survey responses were gathered from workshop participants before, immediately after, and four-months following the workshop. The survey instruments can be found in **Supplemental Index.** One-sided Mann-Whitney U tests [35] were used to compare pre- and post-workshop responses, where our null hypothesis was that there is no difference between the pre- and post-workshop responses and our alternative hypothesis was that the post-workshop responses were higher than the pre-workshop responses. We evaluated each question with *α* = 0.05.

## Results and Discussion

### Educational Jupyter Notebooks

We developed a series of four Jupyter notebooks covering various aspects of kinetic and constraint-based metabolic modeling and metabolic control analysis. A graphical summary of the different areas of metabolic modeling covered and their relationships is shown in **Figure 1**. In addition to learning the theory behind these methods, learners are exposed to the key concepts for successful applications of flux modeling listed below. We also note in **Table 1** and the lesson plans in the **Supplemental Materials** when an exercise can be used to teach one of these concepts.

1. **Concept 1:** The relationship between the noise and time resolution of experimental data and the confidence one can have in parameter estimates and assumed model architectures.
2. **Concept 2:** The uniqueness and identifiability of flux estimates in FBA and 13C-MFA and their relationship to model complexity.
3. **Concept 3:** The distribution of control over fluxes and concentrations in a network across the reactions of that network.

**Figure 1:**
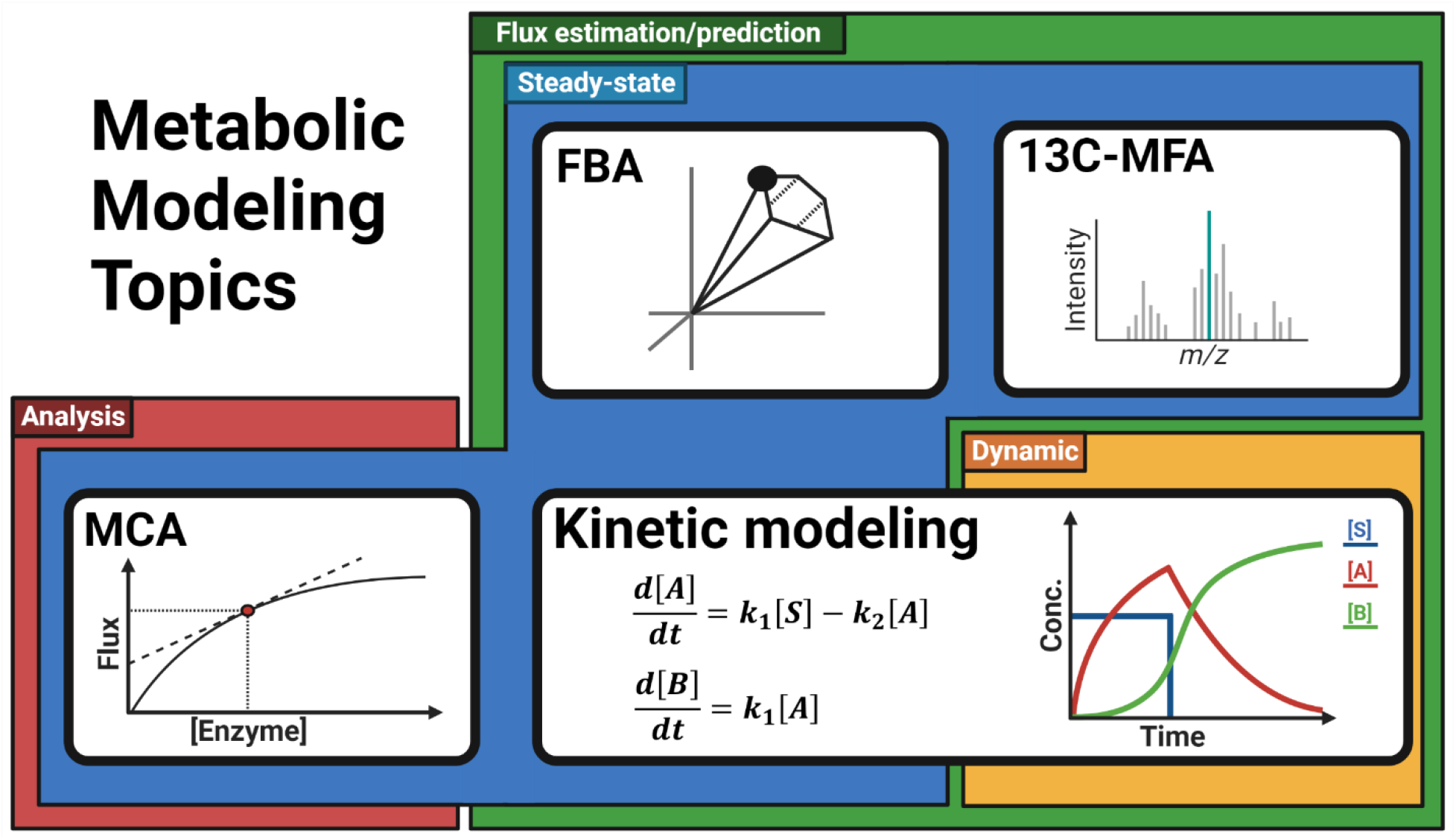
Metabolic modeling topics covered in the resources presented in this study. A majority of the techniques covered – kinetic modeling, FBA, and 13C-MFA – are used to estimate or predict fluxes through a metabolic network. MCA, on the other hand, is used to analyze the relationship between enzyme activities/concentrations and metabolite or regulator concentrations on the flux through the network. Within the flux estimation/prediction techniques, kinetic modeling can be used to estimate fluxes and metabolite concentrations in systems whether they are in steady-state or not (dynamic systems where concentrations are still changing). The constraint-based modeling techniques of FBA and 13C-MFA, on the other hand, rely on an assumption of metabolic steady-state, as does MCA.

These concepts are necessary both to effectively conduct any experiment or study involving flux analysis and to understanding the primary metabolic modeling literature. They are often not intuitively obvious, and the first two also receive rather little attention in the teaching or research literature. The concepts are therefore explained in the lecture notes, revisited throughout the Jupyter notebooks and demonstrated with hands-on exercises. For example, in **Exercises 4.0 – 4.2** in the **Day 4** Jupyter Notebook, learners gain insight into **Concept 1** by first using a kinetic model to generate simulated labeling data and then attempting to fit it using both correctly and incorrectly specified network models using 13C-MFA. By doing so, the learners can observe the difference in 13C-MFA fits when using the correct or incorrect model specification and how this difference can be obscured even by low levels of experimental noise. This allows instructors to highlight important issues concerning data quality and to discuss model selection, which is rarely addressed in the literature [36].

The subjects covered in the sections of each notebook with the timeline for a 4 day workshop are given in **Table 1.** On the first and second days, learners are given an extensive introduction to kinetic modeling theory and exercises before learning about MCA, FBA, and 13C-MFA. We do this to allow learners to gain both a theoretical and practical understanding of the dynamic ways that matter moves through biochemical networks. The hands-on experience exposes learners to the sometimes surprisingly complex behavior of even simple networks governed by systems of Ordinary Differential Equations (ODEs). This is aimed at giving learners a strong sense of the dynamics of metabolic systems before learning about steady-state approaches, in which simplifications of the kinetic state allow powerful analyses in 13C-MFA and FBA. MCA is explored in the second and third days and MCA calculations of flux- and concentration-control coefficients are discussed. Control coefficients are connected to the understanding of reversible first-order kinetics participants gained from the preceding kinetic modeling exercises. Lastly, participants are introduced to constraint-based methods by analyzing the same network structure using kinetic modeling, FBA, and 13C-MFA. This highlights the different inputs needed and the resulting outputs from each technique. To our knowledge, this is the first such cross-comparison of different metabolic modeling techniques presented in the teaching literature, and we believe this will be of value to instructors introducing this material to their students and trainees.

Interactive sliders and drop-down menus were incorporated into all of the notebooks to allow learners to modify parameters, run simulations and visualize their results. This allows learners to expose the underlying simulation code and for those with a modest background in Python or general coding to see how the simulations function and potentially to modify the model structures. By default the code is not visible, making the notebooks approachable for participants interested in using metabolic modeling without engaging with the underlying code. We believe that the incorporation of these interactive modules into the notebooks will make the resources presented in this publication useable by learners with little to no coding knowledge.

In writing the notebooks, special attention was given to commenting the Python code used to run the simulations and interactive interface elements. We believe the extensive commenting used in these notebooks, together with the use of intuitive and easy-to-understand methods for implementing the simulations will make the notebooks both easy for instructors to adopt and for learners interested in the underlying code to understand it. This is in contrast to software like COPASI that, while very powerful, obscure the underlying simulation logic [37]. Installation and compatibility issues are commonplace when using computational resources, particularly when workshop or class participants are asked to run code or software on their own computers. To further ensure maximal useability of these resources by instructors, detailed installation instructions for Windows, MacOS, and Linux systems with the specific version numbers needed to successfully run all of the notebooks provided with the notebooks.

### Implementation in Workshop and Survey Results

The Jupyter notebooks were incorporated into a four-day workshop held at Michigan State University in May 2022. Participants in the workshop included graduate students and postdoctoral researchers. Each day of the workshop consisted of three hours of lecture in the morning and a three-hour hands-on period for computational exercises. Due to time constraints and interest among the participants in constraint-based modeling approaches – particularly label-assisted flux mapping using MFA – the third day’s notebook exercises were omitted and replaced with the fourth day’s exercises on constraint-based modeling. The last day of the workshop was used for an open-ended discussion about participants’ research aims and how they could incorporate what they learned in the workshop into their own work. For instructors interested in incorporating not only the computational resources developed for the workshop, but also all or portions of the lecture material, detailed lecture notes have been provided in the **Supplemental Materials** document.

The pre- and post-workshop survey results suggest that participants felt they gained greater confidence in and knowledge of metabolic modeling over the course of the workshop **(Table 2)**. Our survey evaluated participants’ self-assessed confidence and competence but did not ask participants to attribute their comprehension gains to the lecture or hands-on components. In a free-response question (“What did you find useful about the workshop?”), one participant responded, “Understanding what goes into metabolic modeling, learning how to critically appraise these models in published literature, and beginning to learn how to implement them into our own projects.” In response to that same question, another participant focused more specifically on FBA: “The hands-on use of *cobrapy* was very helpful. This helped me understand how one goes about metabolic modeling.” It should be noted, however, that the sample sizes for the study were small and we had fewer respondents in the post-workshop survey than the pre-workshop survey (N = 12 in the pre-workshop survey and N = 7 in the post-workshop survey). Because of this, the results may be skewed due to survivorship bias from learners who were either no longer interested in the topic or unhappy with the presentation of the material leaving and not participating in the post-workshop survey.

**Table 2:** Quantitative pre- and post-workshop survey results evaluating learners’ self-assessed confidence and competence in metabolic modeling techniques.

| Question | Pre-workshop Median | Post-workshop Median | Significant improvement?<sup>a</sup> |
| --- | --- | --- | --- |
| I feel confident in applying and incorporating metabolic modeling techniques to my research question(s). | 2 | 3 | Not Significant |
| I feel confident in evaluating the results of a metabolic modeling study or a study that incorporates metabolic modeling. | 2 | 3 | Significant |
| I feel confident in identifying metabolic modeling techniques and software that I can apply to my research question(s). | 2 | 4 | Significant |
| I understand the purpose(s) of metabolic modeling. | 4 | 4 | Significant |
| I can describe kinetic metabolic modeling, what information it can provide, and its limitations. | 3 | 4 | Significant |
| I can describe Metabolic Flux Analysis, what information it can provide, and its limitations. | 3 | 4 | Significant |
| I can describe Flux Balance Analysis, what information it can provide, and its limitations. | 2 | 4 | Significant |
| I understand the data types I would need to carry out kinetic metabolic modeling. | 2.5 | 4 | Significant |
| I understand the data types I would need to carry out Metabolic Flux Analysis. | 2.5 | 4 | Significant |
| I understand the data types I would need to carry out Flux Balance Analysis. | 2 | 4 | Significant |
| I can name the language(s) or software package(s) I would use to incorporate metabolic modeling into my own research. | 2 | 4 | Significant |
| I can critically evaluate the application and results of metabolic modeling in publications and presentations relevant to my area of research. | 3 | 4 | Significant |

<sup>a</sup>Statistically significant improvement was defined by rejection of the null hypothesis by the one-sided Mann-Whitney U test [35] at $\alpha = 0.05$ .

Multiple respondents noted that they would have liked to have worked with real datasets in the exercises rather than simulated ones. Given the modifiability and extensive annotation of the notebooks provided, we encourage instructors using the provided resources to add analyses of real datasets that are relevant to their specific audience. We believe this will help provide real-world context for learners as they carry out the exercises.

Although we have packaged and used the materials presented in the context of an intensive workshop, we believe the materials can be adapted to a variety of teaching circumstances. The Jupyter-based simulations could be used for computer lab sessions in a semester-long course, for example, or used as an interactive demonstration in a lecture setting. With the relevant theory taught beforehand, these resources may also be appropriate for undergraduate learning. As noted, the extensive annotation of the code paired with the easy-to-use graphical interface for the exercises also makes them suitable for both learners with extensive and with no prior knowledge of programming.

## Conclusions

Recognizing the absence of resources for teaching the major areas and techniques of metabolic modeling and flux analysis in an integrative fashion, we have developed a set of resources that should be readily adoptable by instructors, students, and researchers alike to teach and learn. By emphasizing the legibility and cross-platform useability of our code, we hope the resources presented in this study can be used and incorporated by the broader teaching community into other workshop and class settings.

## Supporting information

Supplemental Lecture Notes Day 1

Supplemental Lecture Notes Day 2

Supplemental Lecture Notes Day 3

Supplemental Lesson Plan Day 1

Supplemental Lesson Plan Day 2

Supplemental Lesson Plan Day 3

Supplemental Lesson Plan Day 4

Supplemental Survey Instruments

## Data and Code Availability Statement

All code and documentation developed for the present study can be found at https://github.com/Gibberella/Metabolic-Modeling-Lessons.

## Acknowledgements

This research was supported by the Office of Science (BER), U.S. Department of Energy, Grant no DE-SC0018269 (J.A.M.K., A.G., Y.S-H.). This work is supported, in part, by the NSF Research Traineeship Program (Grant DGE-1828149) to J.A.M.K. This publication was also made possible by a predoctoral training award to J.A.M.K. from Grant T32-GM110523 from National Institute of General Medical Sciences (NIGMS) of the NIH. Its contents are solely the responsibility of the authors and do not necessarily represent the official views of the NIGMS or NIH. We would like to acknowledge the Pathways to Research program at Michigan State University for its support of A.G. We would also like to acknowledge Veronica Greve for her input on the survey component of this study.

## Supporting Information

Additional supporting information can be found online in the Supporting Information section at the end of this article.

## References

1 J. Nielsen It Is All about Metabolic Fluxes. (2003) J. Bacteriol. 185, 7031–7035.

2 P. A. Saa, L. K. Nielsen Formulation, construction and analysis of kinetic models of metabolism: A review of modelling frameworks. (2017) Biotechnol. Adv. 35, 981–1003.

3 J. D. Orth, I. Thiele, B. O. Palsson What is flux balance analysis? (2010) Nat. Biotechnol. 28, 245–248.

4 M. R. Antoniewicz Methods and advances in metabolic flux analysis: a mini-review. (2015) J. Ind. Microbiol. Biotechnol. 42, 317–325.

5 M. A. G. Koffas, G. Y. Jung, G. Stephanopoulos Engineering metabolism and product formation in Corynebacterium glutamicum by coordinated gene overexpression. (2003) Metab. Eng. 5, 32–41.

6 M. Koffas, G. Stephanopoulos Strain improvement by metabolic engineering: Lysine production as a case study for systems biology. (2005) Curr. Opin. Biotechnol. 16, 361–366.

7 J. Becker, O. Zelder, S. Häfner, H. Schröder, C. Wittmann From zero to hero-Design-based systems metabolic engineering of Corynebacterium glutamicum for l-lysine production. (2011) Metab. Eng. 13, 159–168.

8 K. H. Lee, J. H. Park, T. Y. Kim, H. U. Kim, S. Y. Lee Systems metabolic engineering of Escherichia coli for L-threonine production. (2007) Mol. Syst. Biol.

9 J. H. Park, K. H. Lee, T. Y. Kim, S. Y. Lee Metabolic engineering of Escherichia coli for the production of L-valine based on transcriptome analysis and in silico gene knockout simulation. (2007) Proc. Natl. Acad. Sci. U. S. A. 104, 7797–7802.

10 S. Gleizer, R. Ben-Nissan, Y. M. Bar-On, N. Antonovsky, E. Noor, Y. Zohar, et al. Conversion of Escherichia coli to Generate All Biomass Carbon from CO2. (2019) Cell. 179, 1255–1263.e12.

11 A. P. Burgard, P. Pharkya, C. D. Maranas OptKnock: A Bilevel Programming Framework for Identifying Gene Knockout Strategies for Microbial Strain Optimization. (2003) Biotechnol. Bioeng. 84, 647–657.

12 D. A. Fell Metabolic control analysis: A survey of its theoretical and experimental development. (1992) Biochem. J. 286, 313–330.

13 R. Moreno-Sánchez, E. Saavedra, S. Rodríguez-Enríquez, V. Olín-Sandoval Metabolic Control Analysis: A tool for designing strategies to manipulate metabolic pathways. (2008) J. Biomed. Biotechnol. 2008,.

14 R. P. Armando, S. J. Francisca, M. Á. Medina First steps in computational systems biology: A practical session in metabolic modeling and simulation. (2009) Biochem. Mol. Biol. Educ. 37, 178–181.

15 G. L. Chaves, R. S. Batista, J. de Sousa Cunha, D. L. Altmann, A. J. da Silva Teaching cellular metabolism using metabolic model simulations. (2022) Educ. Chem. Eng. 38, 97–109.

16 K. W. Wong, J. P. Barford, J. F. Porter Understanding the practical consequences of metabolic interactions - A software package for teaching and research. (2004) IFAC Proc. Vol. 37, 315–320.

17 K. W. W. Wong, J. P. Barford Metstoich: Teaching quantitative metabolism and energetics in biochemical engineering. (2010) Chem. Eng. Educ. 44, 147–156.

18 J. L. Snoep, P. Mendes, H. V Westerhoff Teaching Metabolic Control Analysis and kinetic modelling: Towards a portable teaching module. (1999) Biochem. (Lond). 25–28.

19 C. Rodríguez-Caso, F. Sánchez-Jiménez, M. Á. Medina A modeling and simulation approach to the study of metabolic control analysis. (2002) Biochem. Mol. Biol. Educ. 30, 169–171.

20 C. R. Angelani, P. Carabias, K. M. Cruz, J. M. Delfino, M. de Sautu, M. V. Espelt, et al. A metabolic control analysis approach to introduce the study of systems in biochemistry: the glycolytic pathway in the red blood cell. (2018) Biochem. Mol. Biol. Educ. 46, 502–515.

21 M. R. Antoniewicz A guide to 13C metabolic flux analysis for the cancer biologist. (2018) Exp. Mol. Med. 50,.

22 S. B. Crown, W. S. Ahn, M. R. Antoniewicz Rational design of 13C-labeling experiments for metabolic flux analysis in mammalian cells. (2012) BMC Syst. Biol. 6 43.

23 M. Dieuaide-Noubhani, A. P. Alonso (2014) Plant metabolic flux analysis. Springer,.

24 J. O. Krömer, L. K. Nielsen, L. M. Blank Metabolic Flux Analysis. (2014) Methods Mol. Biol. New York, NY.

25 R. G. Ratcliffe, Y. Shachar-Hill Measuring multiple fluxes through plant metabolic networks. (2006) Plant J. 45, 490–511.

26 G. Stephanopoulos, A. A. Aristidou, J. Nielsen Metabolic engineering: principles and methodologies. (1998).

27 T. Kluyver, B. Ragan-Kelley, F. Pérez, B. Granger, M. Bussonnier, J. Frederic, et al. Jupyter Notebooks—a publishing format for reproducible computational workflows. (2016) Position. Power Acad. Publ. Play. Agents Agendas - Proc. 20th Int. Conf. Electron. Publ. ELPUB 2016. 8790.

28 C. R. Harris, K. J. Millman, S. J. van der Walt, R. Gommers, P. Virtanen, D. Cournapeau, et al. Array programming with NumPy. (2020) Nature. 585, 357–362.

29 P. Virtanen, R. Gommers, T. E. Oliphant, M. Haberland, T. Reddy, D. Cournapeau, et al. SciPy 1.0: fundamental algorithms for scientific computing in Python. (2020) Nat. Methods. 17 261–272.

30 F. Matsuda, K. Maeda, T. Taniguchi, Y. Kondo, F. Yatabe, N. Okahashi, et al. mfapy: An opensource Python package for 13C-based metabolic flux analysis. (2021) Metab. Eng. Commun. 13 e00177.

31 A. Ebrahim, J. A. Lerman, B. O. Palsson, D. R. Hyduke COBRApy: COnstraints-Based Reconstruction and Analysis for Python. (2013) BMC Syst. Biol. 7,.

32 J. D. Orth, T. M. Conrad, J. Na, J. A. Lerman, H. Nam, A. M. Feist, et al. A comprehensive genome-scale reconstruction of Escherichia coli metabolism–2011. (2011) Mol. Syst. Biol. 7, 1–9.

33 J. D. Orth, R. M. T. Fleming, B. Ø. Palsson Reconstruction and Use of Microbial Metabolic Networks: the Core Escherichia coli Metabolic Model as an Educational Guide. (2010) EcoSal Plus. 4,.

34 R. Likert A technique for the measurement of attitudes. (1932) Arch. Psychol. 140, 1–55.

35 M. Neuhäuser in M. Lovric (Ed.) (2011) Springer Berlin Heidelberg, Berlin, Heidelberg, pp. 1656–1658.

36 N. Sundqvist, N. Grankvist, J. Watrous, J. Mohit, R. Nilsson, G. Cedersund Validation-based model selection for 13C metabolic flux analysis with uncertain measurement errors. (2022) PLOS Comput. Biol. 18 e1009999.

37 S. Hoops, R. Gauges, C. Lee, J. Pahle, N. Simus, M. Singhal, et al. COPASI - A COmplex PAthway SImulator. (2006) Bioinformatics. 22, 3067–3074.

