## Supplemental Lecture Notes Day 1 for "Integrative Teaching of Metabolic Modeling and Flux Analysis with Interactive Python Modules"

**Lecture Notes Day 01**

**Pre-reading assignment:**

R. G. Ratcliffe, Y. Shachar-Hill. Measuring multiple fluxes through plant metabolic networks. (2006) Plant J. 45, 490–511.

**Network Biology**

1. Biology operates at the network level, but we typically think about and investigate it in terms of individual pieces or pathways. Whether it’s metabolism, signaling, development, or other processes, very few of them operate meaningfully in the form of linear pathways.
   a. **Examples:**
      i. There is no intermediate in the glycolytic pathway that is not also part of another pathway.
      ii. In the literature on hormonal signaling, there is much discussion of “cross-talk” between hormonal signaling pathways. In reality, this language just refers to the fact that these signaling pathways are found in the context of a larger network.
   b. However, this does not mean that the term “pathway” is meaningless. Rather, it can be a useful way of identifying important operational units within a network.
      i. **Example:** In a human muscle cell that is working hard, the vast majority of flux really will be from hexose to CO2 via the operation of glycolysis and the TCA cycle.
2. Metabolism is just one class of functional biological networks. It is also a very well-studied. The modeling covered in this workshop will concern itself primarily with metabolism, but many of the ideas and principles that go into analyzing metabolic networks may be applied to other biological networks.
   a. Some of the same mathematics can be used to describe the flows, or fluxes, of individuals in ecological research, or the flow of genes in population genetics.

**Mechanistic Modeling in Biology**

1. Much of the content of this workshop will involve analyzing the kinetics models in which the flow of matter, energy, or information is described by relatively simple systems of differential equations.
2. We are at a stage in biology where we have the ability to generate massive quantities of data related to, for example, organisms’ genotypes, transcriptomes, and proteomes. To-date, much of the analysis that has been done on these data types has been statistical in nature, as opposed to functional or mechanistic analyses. In contrast, the form of metabolic modeling we’re going to be doing in this workshop is based on or hypothesizes an explicit structure and mechanistic relationships between the components of a network (i.e. the metabolites involved and their compartmentation, as well as the reactions that interconvert these metabolites).

**Kinetic flux analysis introduction**

We start our investigation of flux modeling with kinetic flux analysis. Kinetic flux analysis involves using time courses to analyze the flow of materials through biological networks. This is done for three reasons:

1. It is historically the oldest of the major metabolic modeling approaches.
2. It is the method that teaches the most about non-metabolic biological network modeling.
3. It is the most intuitive way to think about biological dynamics and learning about it gives insight into how other methods, namely steady-state flux analysis, work. These steady-state methods can be thought of as a subset of kinetic approaches that apply when certain conditions (namely metabolic steady-state) are satisfied.

**Experimental analysis of the fluxes and the properties of metabolic networks**

*Principles and tools of kinetic (dynamic) labeling*

This section references the following publication: [1]

We can consider a simple network as below. A labeled substrate S with a fractional enrichment *f_s_* is introduced into a linear pathway:

$$\begin{aligned} S\underset{\to}{J_{sa}}A\underset{\to}{J_{ab}}B\underset{\to}{J_{bp}}P \#\left( M1 \right) \end{aligned}$$

Where the flow from the substrate (S) to the first intermediate (A) is given by J_sa_, the flow from the first to the second intermediate (B) is given by J_ab_, and the flow from the second intermediate to the product (P) is given by J_bp_. Note that the letter J is often used now to denote fluxes, though historically V is also common. In either case, the subscripts following the letter J or V are used to denote the substrate and product involved in that flow or flux.

For the purposes of this example we’ll assume S is constant. The rates of change of the concentrations of the metabolites A, B, and P, will simply be the rates of influx minus the rates of efflux:

$$\begin{aligned} \frac{d\left[ A \right]}{dt}=J_{sa}-J_{ab} \#\left( E1 \right) \end{aligned}$$

$$\begin{aligned} \frac{d\left[ B \right]}{dt}=J_{ab}-J_{bp}\#\left( E2 \right) \end{aligned}$$

$$\begin{aligned} \frac{d\left[ P \right]}{dt}=J_{bp} \#\left( E3 \right) \end{aligned}$$

If we were to assume metabolic steady-state – that is, that the concentrations of the metabolites remain constant over time, this would mean that the fluxes in and out of A and B would need to equal one another. Therefore:

$$\begin{aligned} J_{sa}=J_{ab}=J_{bp}\#\left( E4 \right) \end{aligned}$$

In this case, instead of three independently varying flux values, we have one. Another way of putting this is that the imposition of steady-state results in us only working with one degree of freedom. In kinetic analysis, in contrast to steady-state analyses, we do not make this assumption, resulting in more degrees of freedom. This greater number of degrees of freedom makes kinetic modeling more computationally and mathematically demanding than steady-state analyses.

The time-dependence of labeling (radioactive or stable isotope) is given by:

$$\begin{aligned} \frac{df_{A}\left[ A \right]}{dt}=f_{s}J_{sa}-f_{a}J_{ab} \#\left( E5 \right) \end{aligned}$$

Where f_n_ is the proportion of metabolite n that is isotopically labeled. In real-world systems, the concentration dependence of each of our fluxes J_sa_ through J_bp_ depends on an underlying catalytic mechanism. For example, under simple Michalis-Menten kinetics:

$$\begin{aligned} J_{sa}=\frac{Vmax\left[ S \right]}{Km+\left[ S \right]}\#\left( E6 \right) \end{aligned}$$

The full set of equations describing the time-dependence of the concentrations and labeling of S, A, B, and P can be used to simulate the response of the system to any change in substrate supply.

**At this point, a walkthrough is given of:**

1. A plot of concentration over time for the model **M1** (from **Fig. B2** in [1]).
2. A plot of fluxes over time for the model **M1** (from **Fig. B3** in [1]).
3. A plot of labeling over time for the model **M1** (from **Fig. B4** in [1]).

**Making Measurements of Labeling**

**Mass spectrometry**

Historically, the oldest method of labeling is radioactive isotopic labeling. But some of the most detailed information can be gained using stable isotopes. Stable isotopes have the same chemistry but a different mass. Examples used through the rest of the workshop will primarily focus on stable isotopic labeling, as radioactive labeling is becoming increasingly rare in flux analysis.

Mass spectrometry is used to detect the molecular mass of intact molecules or fragments. This is very useful for measuring labeling since the incorporate of stable isotopes into molecules will change their mass. This gives sensitive measurements of the identities and levels of many (up to hundreds at a time) metabolites as well as the quantification of stable isotopic labeling levels. Of course, this does require the extraction and frequently also the chemical derivatization of analytes.

Gas chromatographic mass spec (GC-MS) is the type most commonly used in published metabolic flux analyses.

- **At this point, a graphical walkthrough is given of how to relate the ^13^C labeling of a two-carbon molecule to mass spectrometry measurements.**

Amino acids are some of the most common targets for labeling analyses because they are (1) found in large amounts, (2) they share common functional groups and so can be derivatized and analyzed as a group, and (3) they are made directly or indirectly from important central metabolic intermediates, giving a readout of labeling across central metabolism.

**Nuclear Magnetic Resonance (NMR) spectroscopy**

Less used, but at least as informative in principle, is NMR. NMR detects signals from nuclear magnetism of naturally abundant isotopes (e.g. 1H, 31P) and of stable isotopes present in low abundance (e.g. 13C, 15N, 2H).

Gives measures of the identities and levels of the more abundant metabolites (>10 micromolar), so it’s not as sensitive a technique as MS. However, it allows quantification of stable isotopic labeling levels without standards and can be used on living samples (*in vivo*) or after extraction in a non-destructive way, allowing you to reuse samples for other analyses.

- **At this point, a graphical walkthrough is given for interpreting an NMR spectrum of a ^13^C labeled glucose molecule, with an emphasis on how NMR is able to distinguish labeling in the different positions of the molecule and the information we gain from this.**

**How easy is it to get data from different labeling methods? What are the pros and cons?:**

1. Radioactive labeling: You have the advantage of enormous sensitivity and ease of detection. But, all ^14^C-labeled molecular species and give the same signal, so you need to separate compounds completely to get useful data. Molecules will give no radioactive signal unless they’re labeled, so metabolite levels are not measured.
2. For mass spectrometry, you get a series of spectra by taking a series of samples, each of which must be isolated, quenched, extracted, frequently derivatized, run on the mass spectrometer, and then the spectra analyzed, with the analysis often taking the most time. This makes the method quite time and labor intensive, though highly informative.

**An example of an *in vivo* NMR time-course experiment and model fitting**

**Note:** This example is taken from unpublished work done in the Shachar-Hill lab [2]

**Experimental setup:** NMR was used to measure the signal from D_2_O inside and outside a plant root. To distinguish between the water inside and outside of the root, an inert paramagnetic agent was added to the area outside the root to shift the signal from water.

The area of the internal peak in successive deuterium NMR spectra from this experiment looks like this:

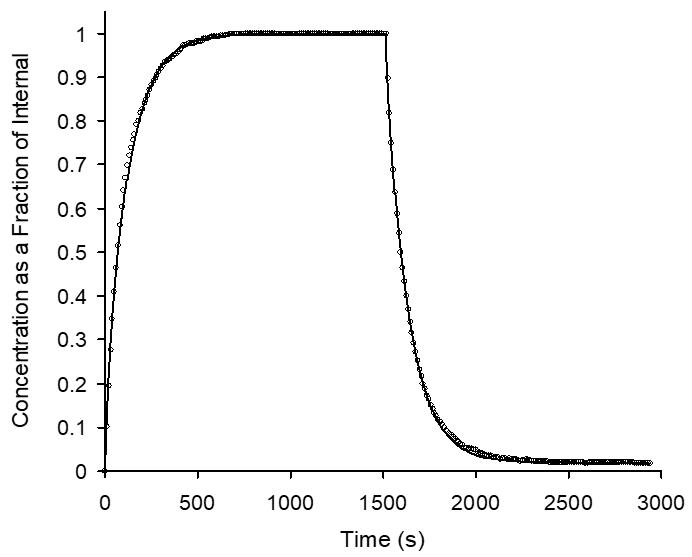
Where the x-axis is time in seconds and the y-axis is the fractional labeling of the water on the inside of the root. The dots represent actual datapoints starting with the addition of D2O to the medium followed after steady state labeling is reached by a washout or chase period after the medium is replaced with unlabeled medium. The line represents the predicted fractional labeling over time given a kinetic model of the system. These predicted values require a model and the model requires a mechanistic description of how the system is operating. In this case, there are multiple mechanistic models that could be used to describe this system, summarized in the image below.

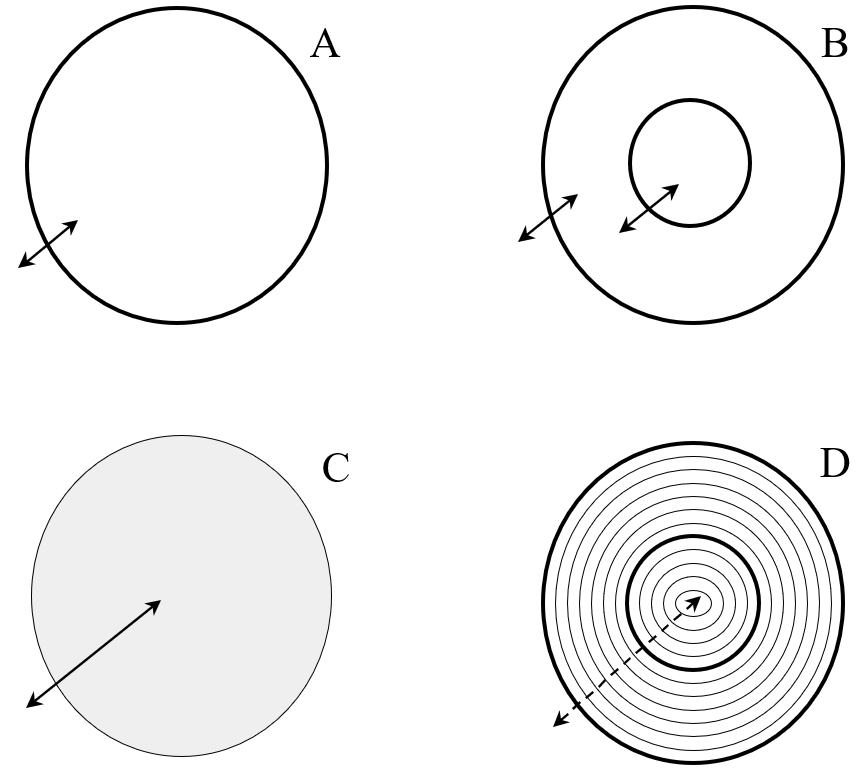

1. In model A, all water inside the root and all water outside the root is treated as the same, or as occupying the same homogeneous compartment. There is one fitted parameter: the rate constant describing movement across the single barrier to water entry.
2. In B, we add an additional barrier that must be crossed (representing the root endodermis), separating the system into three compartments. There are two fitted parameters, corresponding to the rate constant of movement across the first barrier and a separate rate constant for movement across the second.
3. In C, the barrier to water movement is continuous and there is one parameter: the diffusion constant of water inside the root.
4. Finally, in D, there are a large number of small barriers, corresponding to the number of cell layers observed in an actual root under microscopy, as well as an endodermis barrier. There are three parameters in this model. The first and second are the rate constants for crossing the outer and endodermis barrier. The third is the rate constant for crossing each of the cell layers .

Each of these representations of the network can be turned into a set of equations. We can then get best-fit parameters for each of these models and evaluate how well the model predictions line up with our observations.

Going from A to D, we get progressively better fits, as we see in the image below.

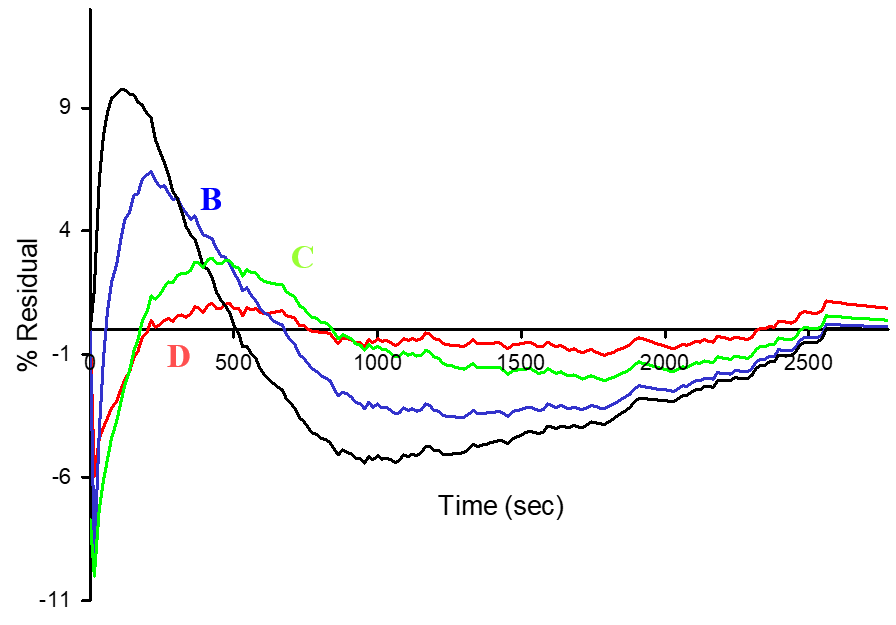

Note that, in general, adding more parameters should allow us to get a better fit, and if more complex models contain simpler models in full (i.e. the models are nested) then the more complex models will always have at least as good a fit as the simpler models. At a certain point, the addition of more parameters or complexity to a model becomes statistically unjustified and one can test whether the addition of a parameter to a model is “worth it” statistically using techniques from the area of model selection. In practice this is (regrettably) rarely done in the realm of metabolic modeling and will not be covered in detail in this workshop.

**To summarize:**

1. The quality of your data determine how refined a model you can construct of a system.
2. You can test hypotheses by building different models that represent something about the structure and relative weights of different aspects of your system and observe how well the model predictions agree with the experimental data.

**Overview of strategy for kinetic analysis of metabolic networks**

1. Start with a real-world system. That is, we start with a real biological network.
2. Generate experimental time course data (usually isotopic labeling measurements) as a readout of the behavior of this network.
3. Build a model, or set of models, of this network out of differential equations to represent the steps or processes in the model.
4. Come up with best-fit values of the model parameters by fitting the model(s) output from Step 3 to the measurements from step 2.
5. Use these parameters to better understand the regulation of the network and make predictions.

And just as a reminder that this sort of kinetic analysis need not be limited to metabolic networks, we can see **Figure 8.3** from [3] for an example of applying this same kind of workflow in the context of gene expression layered on top of a metabolic process, where we can see the interplay of rates of transcription, translation, and enzymatic activity.

**Computational limitations**

The more realistic a model you are working with, the more difficult it becomes to retrieve accurate parameter estimates from kinetic modeling (or any modeling). In this study, the authors set the parameters for all the processes and generated *in silico* data from their simulations of metabolite, transcript, and protein levels at steady-state. They then fed their model of the system and their *in silico* data back into a fitting algorithm to see if it could arrive at the original parameter estimates that generated the *in silico* observations. What they found was that it was possible to get good parameter estimates back, but that the ability to do so was highly sensitive to the quality of the data (in this case, whether noise had been added).

**A kinetic experiment of choline metabolism**

**Reference:** McNeil,S.D. *et al.* (2000) Metabolic modeling identifies key constraints on an engineered glycine betaine synthesis pathway in tobacco. *Plant Physiol.*, **124**, 153–162.

As an example of this process, in this study a labeled substrate was fed to tobacco leaf tissues and the label’s path through a metabolic network was traced to see which, or what combination, of pathways were used to produce a product of interest (phosphatidylcholine, or PC) [4]. By measuring and simulating the labeling of the metabolites involved in the network and using a simple Visual Basic interface of a similar structure to the interactive Python modules in this workshop, the Michaelis-Menten kinetic parameters of the various reactions in this network were fitted. The upshot of this analysis was that overexpression of enzymes catalyzing some processes that had been targeted in earlier work was insufficient to improve production of the metabolite of interest. Competing processes took away too much of the upstream intermediate and the transporter that was responsible for getting the intermediate to the right compartment to be acted on by those overexpressed enzymes was simply not active enough.

The takeaway here is that by building out this kinetic model, researchers were able to ask – quantitatively – what it would take in order to achieve a practical bioengineering goal.

**An example from secondary metabolism – benzenoid fragrance**

**References:** [5,6]

There is a >15 year collaboration among researchers at Purdue (Natalia Dudareva, David Rhodes, and John Morgan) that has combined kinetic metabolic flux analysis with molecular genetics and biochemistry to elucidate many aspects of benzenoid biosynthesis. These benzenoid compounds are fragrances that are emitted from plants and synthesized from a common precursor: phenylalanine. Labeling of this system is done by providing isotopically labeled phenylalanine to flowers and then capturing and analyzing volatilized compounds and some of the intracellular metabolites. We can see a proposed set of pathways constituting the network that generates these volatile compounds in **Figure 1** of [5] and **Figure 1** of [6].

To get a sense of how this system developed, we can see Boatright *et al*. 2004, where first-order rate constants were fitted for the reactions in this network. Notably, no confidence intervals were calculated for any of these reactions. Fast-forward to Marshal Colon *et al.* 2010 and the group had Michaelis-Menten kinetic parameters for each reaction along with 95% confidence intervals. If we refer to **Table 1** in [6] we can see that many of the parameters have massive 95% CIs, meaning that we are not sure what those parameter values really are.

**References**

1	R. G. Ratcliffe, Y. Shachar-Hill Measuring multiple fluxes through plant metabolic networks. (2006) *Plant J.* **45**, 490–511.

2	J. Rosenberg, Y. Shachar-Hill. Unpublished results.

3	P. Mendes in H. Kitano (Ed.) (2001) Found. Syst. Biol., The MIT Press, , pp. 163–186.

4	S. D. McNeil, D. Rhodes, B. L. Russell, M. L. Nuccio, Y. Shachar-Hill, A. D. Hanson Metabolic modeling identifies key constraints on an engineered glycine betaine synthesis pathway in tobacco. (2000) *Plant Physiol.* **124**, 153–162.

5	J. Boatright, F. Negre, X. Chen, C. M. Kish, B. Wood, G. Peel, et al. Understanding in Vivo Benzenoid Metabolism in Petunia Petal Tissue. (2004) *Plant Physiol.* **135**, 1993–2011.

6	A. M. Colón, N. Sengupta, D. Rhodes, N. Dudareva, J. Morgan A kinetic model describes metabolic response to perturbations and distribution of flux control in the benzenoid network of Petunia hybrida. (2010) *Plant J.* **62**, 64–76.
