## Supplemental Lecture Notes Day 2 for "Integrative Teaching of Metabolic Modeling and Flux Analysis with Interactive Python Modules"

**Lecture Notes Day 02**

**Metabolic Flux Analysis to examine production and consumption rates of metabolites in a biological system**

Referenced studies: [1,2]

Labeling studies of the photosynthesizing leaf by introducing CO_2_ incorporating either ^14^C (^14^CO_2_) or ^13^C (^13^CO_2_) to study carbon assimilation and the Calvin-Benson cycle were being done all the way back in the late-40s, so it may seem strange to continue building models of this system. There were several reasons for this, all of which speak to the various motivations researchers have for conducting flux analyses in the first place.

1. There are people who just like doing metabolic flux analysis. It’s what they are good at and they are interested in developing and refining the techniques and that in and of itself is a valuable endeavor to them.
2. After the elucidation of major pathways, unsolved biological questions and unexplained observations related to the labeling patterns of central carbon metabolic intermediates remained.
3. There are those who have suspicions about how *in vitro* investigations translate to what a biological system is doing in its natural state *in vivo*. In this case (and for other systems), metabolic modeling using in vitro enzyme kinetic parameters has produced predicted flux maps, but other researchers believe that these need to be tested with data obtained *in vivo*.
4. In this specific case, there were aspects of some of the previous MFA studies of central carbon metabolism in photosynthesizing leaves that were incomplete or appeared to be based on biologically problematic assumptions, motivating a revisiting of the general area. Successive studies to improve the fidelity and scope of metabolic modeling were discussed in lecture 1.

*An overview of the steps taken in this, and any, 13C-MFA study*

1. Experimental design
2. Labeling experiment
3. Data collection (here LC/GC-MS)
4. If doing instationary 13C-MFA, repeat step 3 across an entire time course. For steady state MFA just take endpoint measurements
5. Using a suitable software package (e.g. INCA [3]), build a metabolic model, typically based on past biochemical literature.
6. Estimate fluxes through the metabolic network architecture from step 5 by optimizing the agreement between model-simulated values and the input data from 3-4
7. Statistically analyze the resulting flux map and uncertainties in flux estimates to determine whether the fit is acceptable and the information sought out in the experiment has been sufficiently well captured

**Studies vary in how physiologically relevant their flux maps are**

In the case of [1,2], the experiments described were done on leaves attached to the rest of the plant and growing under controlled laboratory conditions. Although these are different from, say, field conditions, the gas exchange measurements are made *in vivo* and the sample extracts that are then analyzed by MS are from leaves quenched very rapidly so as to stop enzymatic activity and capture, as best as possible, the levels and labeling of metabolites in the photosynthesizing plant. In other cases, like the benzenoid studies referenced in the Day 1 lecture [4,5], the system is less physiologically relevant. In those studies, flowers needed to be removed from plants and fed very high concentrations of phenylalanine in order to make the necessary measurements, which is substantially different from real-life conditions. This does not make the experiments invalid, as the authors were able to identify and then independently validate a number of features of the system revealed using these techniques, but it does highlight the importance of considering, when one is conducting a flux mapping study or reading about one in the literature, what level of physiological relevance is needed to answer one’s questions.

For pathway discovery in particular, less physiologically relevant systems like the one described in [4,5] are very acceptable, as it’s unlikely a flower will change the pathway it employs to produce a secondary metabolite because it has been removed from a plant. It is likely, however, that it will produce different amounts or proportions of those metabolites – therefore, if these are the questions that one was interested in asking, considering the physiological relevance of the system would be key.

**Details of the study**

1. Gas exchange measurements were made using a LICOR instrument.
2. Each measurement was taken from a single leaf of a *Camelina sativa* plant. The leaf is rapidly quenched to prevent further reactions and then removed from the plant.
   a. Because of how fast some of the initial reactions in photosynthesis occur, an extremely fast quenching setup was used.
   b. Removal of a leaf could perturb the metabolism of the rest of the plant, so each leaf used for a measurement is taken from an independent plant.
3. Sample extracts were divided, prepared for analysis, and run on reverse phase LC/MS/MS, GC/MS, and ion exchange LC/MS/MS instruments.
4. Three replicates of each sample were taken for each timepoint, with timepoints going out from 0 to 60 minutes.
5. Interpretation of data:
   a. **Terminology:**
      i. Isotopologues are molecules that differ only in their isotopic composition. For example, glycine molecules with zero, one, or two ^13^C atoms incorporated are isopologues.
      ii. Isotopomers are molecules that differ not in their isotopic composition, but in the position of the isotopes. So, glycine molecules with one ^13^C incorporated (M+1) could be labeled at either position 1 or 2, and these would be isotopomers.
   b. The fitting process involves a software package (in this case [3]) starting with a random flux map and using this to calculate, given the known label provided, setup, and other measured experimental constraints, what the labeling pattern should be for all the intermediates in the network. This prediction is compared to the actual measured labeling data and the flux map is iteratively improved from there until an optimal fit is arrived at. The goodness of fit is statistically tested for acceptability.
   c. **Compartmentation**: One difficulty is that in eukaryotic central metabolism, multiple key metabolites are located in more than one cellular compartment, but the extraction process will simply give you all of, for example, the glycine in a sample. To counteract the uncertainty this introduces, one must make more measurements of more intermediates in metabolism in order to have enough information to make confident flux estimates.
   d. **Inability to get good signal for some metabolites:** Due to either low concentration, instability, or the analytical methods used, you will not be able to get usable measurements of all metabolic intermediates. Oxaloacetate and α-ketoglutarate are two examples from the TCA cycle, but there are many others. Once again, this also requires us to make many more measurements and/or approximations/assumptions to compensate for the lack of direct experimental information.

**Results of study** [1]

During the day, some of the CO2 assimilated in photosynthesis is reemitted. Some of this emission can be attributed to the process of photorespiration, but the remainder that cannot is called R_L_, or respiration in the light. The flux map resulting from this study was used to figure out the source of R_L_. The operation of an oxidative pentose phosphate (OPP) shunt was found to be the major source of this daytime respiration, rather than the TCA cycle, which had previously been proposed as the source.

Flux analysis papers are one-off studies with no substantial follow-up. But, in this case, another study was done [2] which added on additional components to the metabolic network whose existence was suggested by a statistical model selection exercise.

**Evaluating flux analysis studies from the literature, or “How skeptical should I be?”**

- When fitting a line or other model to a set of data, one typically has more datapoints than are necessary to fit the model parameters (i.e. the model is overdetermined). This requires using a regression approach to come up with the best-fit parameters given that the model predictions will not hit each data point exactly.
- Different parameters will have different sensitivity to different datapoints. This must be considered at the experimental design and interpretation stage. Another way of thinking of this is that not all data are equally important. Even if you have a lot of data, you may be unable to determine some or all parameters sufficiently.
- In flux maps like those presented in [1,2], almost all the data is not direct flux measurements. Indeed, the only directly flux measurements are “external” fluxes of metabolites entering the system as substrates or leaving as products (here gas exchanges and the starch synthesis rate). This makes flux measurements seem to many biologists – rightly so – like they may be less reliable, due to the fact that they estimated indirectly from labeling rather than directly measured. The fitted fluxes from a set of labeling data using an incorrectly specified model may look acceptable but be inaccurate.
  - For this reason, it is appropriate to see whether the model has been tested by direct measurement of some fluxes to validate the predictions, by comparing some global features of the flux map to independent measures of these features, or with additional non-redundant labeling data.

**Computational aspects:**

**Finding the global best fit(s)**

When using any linear or non-linear regression-based approach, one must take measures to ensure that the best-fit set of parameters is, in fact, the global best fit and not simply a local minimum. To ensure this:

1. One should use a robust optimization algorithm. Luckily, such methods are built into software packages like [3].
2. One should use many random starting points for their regression.

**Calculating confidence intervals**

Estimating confidence intervals in flux analysis is actually quite computationally challenging, but a few different methods exist, including local sensitivity analysis, parameter continuation, and Monte Carlo sampling. Local sensitivity analysis turns out to give inaccurate confidence intervals, so parameter continuation and Monte Carlo sampling are the methods now commonly used. It is our opinion that Monte Carlo sampling is the gold-standard method, but parameter continuation is increasingly the default method implemented in software packages like INCA due to its favorable computational characteristics.

Monte Carlo sampling involves randomly altering the input datapoints based on their observed experimental measurement variabilities and re-fitting the model. By doing this many times, one can generate 95% confidence intervals for all of the fitted parameters (i.e. fluxes) in the network and get an estimate of which fluxes – net, forward, and reverse – are more confidently estimated and which are less.

**Summary of questions one should ask:**

1. Is the definition of the metabolic network justified?
2. What is the quality of the data?
3. Is the complexity of the model justified by the data available?
4. What is the reliability of the parameter estimations?
5. Were independent tests of the model and its conclusions made?

**Metabolic Control Analysis**

**References for this section:** [6,7]

Metabolic Control Analysis (MCA) is a framework for determining how the kinetic behavior of a pathway or network can be explained in terms of the properties of the individual enzymes.

For a single metabolic step, the flux is simply dependent on the kinetic properties and level of the enzyme, the substrates, and regulators/effectors. This is intuitive enough. What’s less intuitive is that:

1. The dependence of flux on enzyme level (or activity) is non-linear.
2. The slope of the relationship between flux and enzyme level (or activity) is different for different enzymes.
3. The slope of the relationship between flux and enzyme level is different under different conditions.

What MCA aims to do is figure out how control over the flux through a metabolic network is distributed across the enzymes catalyzing its reactions. A key observation one makes when doing MCA is that there are no “rate-limiting steps” that is, individual metabolic reactions or enzymes that entirely determine the overall rate of a process. Instead, control over the overall rate is distributed across the network, though some enzymes may, in some cases, have very high degrees of control. Another observation is that the control exerted by an individual enzyme is a function of the activities of all the other enzymes in the network.

MCA can also be used to interrogate how the concentrations of metabolic products or intermediates depend upon enzyme activities, which is very important because these concentrations are often what bioengineers aim to modify.

**Theoretical framework**

**Flux Control Coefficients**

To quantify the control an enzyme has over the flux through a network, we want to calculate the fractional change in that flux over the fractional change in the enzyme concentration.

$$C_{E}^{J}=\frac{\frac{\partial Flux}{Flux}}{\frac{\partial\left[ E \right]}{[E]}}$$

Which we can rewrite as:

$$C_{E}^{J}=\frac{\partial\ln J}{\partial\ln[E]}$$

An enzyme that was a true rate-limiting step would have a flux control coefficient of 1 and one with absolutely no control would have a flux control coefficient of 0. As a reminder, what we actually see in metabolism is a distribution of flux control coefficients between 0 and 1 across all the enzymes in a network. It is useful to note that in a network the sum of all the flux control coefficients will always equal 1.

$$\sum_{i=1}^{n} C_{E_{i}}^{J}=1$$

**Concentration Control Coefficients**

We can also ask how the concentrations of intermediate metabolites are controlled by different enzyme activities. Similar to above, if we are looking the concentration of an intermediate A:

$$C_{v}^{[A]}=\frac{\partial\ln[A]}{\partial\ln v}$$

Where *v* is an enzyme activity and [A] is the concentration of an intermediate A. There is an analogous, but slightly different, summation theorem:

$$\sum_{i=1}^{n} C_{v_{i}}^{[A]}=0$$

**Elasticities**

We can also calculate elasticities, which are local parameters that describe the dependence of the rate of a *particular* reaction in the network on its substrate and product concentrations. Note that in this case, because we’re examining the behavior of a single reaction rate J_i_, rather than the overall flow through the system (J), this local quantity, elasticity, does not depend on the behavior of the rest of the network.

$$E_{ij}=\frac{dJ_{i}}{d\left[ S_{j} \right]}$$

**Examples of MCA from the literature**

*S. cerevisiae* tryptophan biosynthesis

Reference: [8]

In this study, MCA was employed to identify a strategy for improving the flux towards tryptophan biosynthesis in *S. cerevisiae.* The authors were able to over- and under-express enzymes involved in tryptophan biosynthesis. This allowed them to measure, for each enzyme, the relationship between its activity, measured *in vitro* after extraction from the cells, and the rate of tryptophan biosynthesis in those cells. For data, see **Figure 1** and **Table 2** of [8]**.** An interesting thing to note in this study is that, in accordance with the predictions of MCA, obtaining very large increases in overall flux towards tryptophan requires upregulation of a substantial fraction (or all) of the enzymes involved in producing tryptophan.

*Aspergillus*-based cell wall degradation with increased conversion of cell wall sugars to citric acid as well as increased concentration of arabinitol

Reference: [9]

In contrast to the tryptophan study, this one involved building a kinetic model by measuring enzyme kinetics for the enzymes involved in the target pathway, enzyme concentrations, and the concentrations of metabolites and co-substrates. This kinetic model was then used to perform the MCA predictively, by computing the effects of varying enzyme levels on flux. The result of this analysis was that the level of arabinose substrate going into the system had a big effect on the control coefficients and that increasing the concentrations of specifically the first three enzymes in the pathway would be necessary to increase flux through this pathway.

**References**

1	Y. Xu, X. Fu, T. D. Sharkey, Y. Shachar-Hill, and B. J. Walker The metabolic origins of non-photorespiratory CO2 release during photosynthesis: a metabolic flux analysis. (2021) *Plant Physiol.* 1–18.

2	Y. Xu, T. Wieloch, J. A. M. Kaste, Y. Shachar-Hill, T. D. Sharkey Reimport of carbon from cytosolic and vacuolar sugar pools into the Calvin-Benson cycle explains photosynthesis labeling anomalies. (2022) *Proc. Natl. Acad. Sci.* **119**, e2121531119.

3	J. D. Young INCA: A computational platform for isotopically non-stationary metabolic flux analysis. (2014) *Bioinformatics*. **30**, 1333–1335.

4	J. Boatright, F. Negre, X. Chen, C. M. Kish, B. Wood, G. Peel, et al. Understanding in Vivo Benzenoid Metabolism in Petunia Petal Tissue. (2004) *Plant Physiol.* **135**, 1993–2011.

5	A. M. Colón, N. Sengupta, D. Rhodes, N. Dudareva, J. Morgan A kinetic model describes metabolic response to perturbations and distribution of flux control in the benzenoid network of Petunia hybrida. (2010) *Plant J.* **62**, 64–76.

6	D. Fell, A. Cornish-Bowden (1997) Understanding the control of metabolism. Portland press London, .

7	A. Cornish-Bowden (2013) Fundamentals of enzyme kinetics. John Wiley \& Sons, .

8	P. Niederberger, R. Prasad, G. M. Ii, H. Kacsert A strategy for increasing an in vivo flux by genetic manipulations. The tryptophan system of yeast. (1992) *Biochem. J.* **287**, 473–479.

9	M. J. L. de Groot, W. Prathumpai, J. Visser, G. J. G. Ruijter Metabolic Control Analysis of Aspergillus nigerl-Arabinose Catabolism. (2005) *Biotechnol. Prog.* **21**, 1610–1616.
