## Supplemental Lecture Notes Day 3 for "Integrative Teaching of Metabolic Modeling and Flux Analysis with Interactive Python Modules"

**Lecture Notes Day 03**

**What is steady state?**

Equilibrium refers to a state where, for a particular reaction, the rates of the forward and reverse reactions are the same. This is different from steady state, which refers to a system where the concentrations of all intermediates remain constant.

Note that for the purposes of Metabolic Flux Analysis in particular, we need to further distinguish between two types of steady-state: metabolic and isotopic steady-state.

1. Metabolic steady state means that the concentrations of metabolic intermediates and the fluxes between them are constant.
2. Isotopic steady state means that the labeling pattern of metabolic intermediates is constant.

In today’s examples, we are discussing the theory underlying flux analysis in systems that are in both metabolic and isotopic steady state.

**Utility of steady-state data**

In contrast to kinetic analyses where we’re looking at either concentrations or labeling distributions (or both) vary over time, it seems like looking at steady-state labeling may be less informative, or even completely uninformative. How much information can we get out of steady-state labeling data? The answer is: it depends on the labeling strategy used and the network under consideration.

In the case of the benzenoid studies discussed in Day 2, steady-state labeling of the phenyl ring of the substrate phenylalanine would be uninformative from a flux analysis perspective, because the phenyl ring moves unmodified through metabolism and at steady-state all you’ll see is the exact same labeling pattern in the intermediates and products as you do in the substrate. Note that this information can be used to aid in pathway delineation, but it provides no information about relative rates through alternative routes through the network or their regulation.

A noteworthy and dramatic example of how steady-state labeling can be uninformative is studying carbon assimilation in a photosynthetic organism using ^13^CO2 or ^14^CO2. At steady state, all of the metabolic intermediates (and each carbon position in each intermediate) will become labeled to the same degree as the input labeled CO2, giving you no information whatsoever except for a little information on kinetic isotope discrimination if the substrate is partially labeled.

**Challenges associated with steady state**

One difficulty of empirical studies done on steady-state systems is ensuring that the system being studied actually is in steady-state. It can be difficult to maintain a system in metabolic steady-state and it can be even more difficult to keep it in metabolic steady-state long enough to achieve isotopic steady-state.

In cultured cells, exponential growth is a steady state since the specific growth rate will be constant. Tissues that are not actively growing or senescing are also in steady state. With the exception of chemostat cultures, none of these systems are truly in steady state. The specific growth rate of a cell culture in exponential growth, for example, will not remain perfectly constant due to density-dependent changes in growth. Leaves are changing continuously over their developmental timeline. But many systems can be said to be in pseudo steady-state (where overall changes in the system are sufficiently slow compared to the experimental time course and to the metabolic fluxes within the system) and can be maintained in this state for some time.

Another challenge is the fact that the labeling readouts of an experiment are often measured from abundant compounds like protein, starch, cell wall components, or even sucrose. But due to the abundance of these compounds, there will be a substantial number of unlabeled molecules present from before labeling was started that are diluting out the labeling signal. In order to get around this, in microbial studies measurements will be made at or after five doublings have occurred. With five doublings, only about 3% of the total biomass in the measured sample will come from before the labeling experiment started. This small amount can then be accounted for in calculations. This can be more challenging to achieve with tissues developing in culture, and larger corrections for unlabeled preexisting material may be necessary.

**Why do studies on systems at isotopic steady state?**

In the first day’s lecture, we went over a 13C-MFA study done at metabolic, but not isotopic, steady-state. In the literature, this is commonly referred to as isotopically nonstationary MFA [1], or INST-MFA. What are some of the benefits and reasons for doing flux analysis at isotopic steady state? The reasons are both historical and practical.

1. To do isotopically nonstationary MFA, you need to be able to take many time points rather than just one endpoint measurement.
2. To do isotopically nonstationary MFA, you need to be able to add and/or remove label rapidly relative to the fluxes being investigated. In some systems this will not be feasible.
3. There are fewer compounds to analyze and these compounds can be in greater quantity (e.g. high concentration end products like amino acids in proteins) in isotopic steady-state analysis in comparison to isotopically nonstationary analysis. This reduces the detection sensitivity or tissue quantity necessary for the experimental work.

**Steady-state analysis theory**

To demonstrate the theory behind steady state analysis, we can examine the very simple metabolic network shown below.

**Terminology:**

1. Each circle represents a metabolite. We’ll refer to these as “nodes” or “vertices.”
2. Each arrow represents a reaction. We’ll refer to these as “edges.”

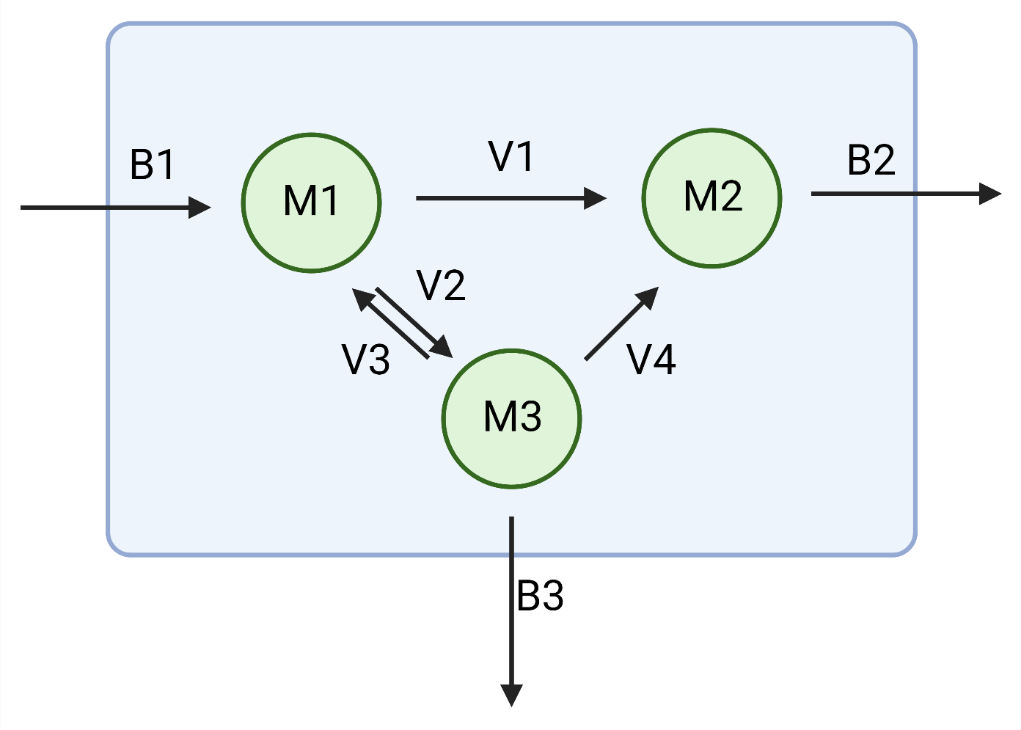

**Model details**

Note that we have two arrows between M1 and M3 representing a reversible reaction and that we have a boundary across which three fluxes, b_1_ through b_3_, cross to enter and leave the network. This boundary could represent a cell or organellar membrane, or it could simply represent an arbitrary delineation between a particular metabolic subnetwork of interest and the rest of metabolism. In either case, these fluxes spanning the boundary are “external fluxes” and all of the fluxes within the boundary are “internal fluxes.”

We are, from this point forward, only dealing with the value of the fluxes rather than the kinetic parameters that ultimately give rise to those fluxes. We are simply treating the fluxes as parameters whose values we are trying to estimate.

**Dynamic mass balance**

This system can, as was done in the first day’s lecture, be represented as a system of ordinary differential equations describing the rates of change of the concentrations of the internal metabolites:

$$\frac{d[M1]}{dt}=-v_{1}-v_{2}+v_{3}+b_{1}$$

$$\frac{d[M2]}{dt}=v_{1}+v_{4}-b_{2}$$

$$\frac{d[M3]}{dt}=v_{2}-v_{3}-v_{4}-b_{3}$$

We can also represent the reactions in this network and how they relate to each metabolite using something called a *stoichiometric matrix* S, where each column represents a reaction and each row a metabolite, with the values at each position representing the stoichiometric coefficient associated with a metabolite in a particular reaction. The stoichiometric coefficient being the number of molecules of a metabolite produced (when positive) or consumed (when negative) in a reaction. For this network …

$$\boldsymbol{S}=\left[ \begin{matrix} -1 & -1 & 1 & 0 & 1 & 0 & 0 \\ 1 & 0 & 0 & 1 & 0 & -1 & 0 \\ 0 & 1 & -1 & -1 & 0 & 0 & -1 \end{matrix} \right]$$

Conveniently, if we multiply this matrix by a vector of all the flux parameters in our network, we get back the equations for the system of differential equations making up our network. That is to say:

$$\left[ \begin{matrix} \frac{d\left[ M1 \right]}{dt} \\ \frac{d[M2]}{dt} \\ \frac{d[M3]}{dt} \end{matrix} \right]=\left[ \begin{matrix} -1 & -1 & 1 & 0 & 1 & 0 & 0 \\ 1 & 0 & 0 & 1 & 0 & -1 & 0 \\ 0 & 1 & -1 & -1 & 0 & 0 & -1 \end{matrix} \right]\left[ \begin{matrix} v_{1} \\ v_{2} \\ v_{3} \\ v_{4} \\ b_{1} \\ b_{2} \\ b_{3} \end{matrix} \right]$$

Recall that at steady-state, the rate of change of the concentration of each metabolite is zero. We can therefore rewrite the equation above as:

$$\left[ \begin{matrix} 0 \\ 0 \\ 0 \end{matrix} \right]=\left[ \begin{matrix} -1 & -1 & 1 & 0 & 1 & 0 & 0 \\ 1 & 0 & 0 & 1 & 0 & -1 & 0 \\ 0 & 1 & -1 & -1 & 0 & 0 & -1 \end{matrix} \right]\left[ \begin{matrix} v_{1} \\ v_{2} \\ v_{3} \\ v_{4} \\ b_{1} \\ b_{2} \\ b_{3} \end{matrix} \right]$$

Which turns a calculus problem into an algebra problem. The solutions to this problem will be all of the vectors of fluxes that, when multiplied by our stoichiometric matrix, give a vector of zeros. That is, all vectors of fluxes that are consistent with the imposition of steady state.

We refer to the imposition of steady state as a **constraint** in this context because it constrains the space of feasible flux vectors that can be returned as solutions. Based on experimental or theoretical grounds, we typically define additional constraints in these kinds of steady-state analyses, including requiring that certain fluxes must be zero or positive, as well as setting maximum possible values. Because the math we’re doing relies on these constraints to delimit the space of possible solutions, MFA and FBA are both referred to as “constraint-based metabolic modeling.”

Observe that the number of reactions in our network exceeds the number of metabolites. In general, this will be the case for metabolic networks. Because of this, the matrix S will have more columns than rows. Because of this, even with the additional constraints we’ve applied, the set of viable flux distributions for the network will be infinite.

At this point, the two main approaches in constraint-based modeling (FBA and MFA) diverge.

1. One way of applying additional constraints is to consider labeling data. **This approach is MFA**.
2. Another way is to apply linear optimization principles to identifying unique or smaller sets of solutions within the full solution space. **This is FBA.**

The optimization done in FBA is with respect to maximizing (or minimizing) the value of an **objective function** Z. The objective function represents a flux or set of fluxes whose value or linear sum respectively, we assume the organism is attempting to maximize or minimize. Expressed mathematically:

$$Z=\sum_{j=1}^{n} c_{j}v_{j}$$

Where *c* is a vector of coefficients (typically 1), *v* is a vector of fluxes, and n is the number of fluxes in the objective function Z. This value will be maximized or minimized, subject to the other constraints on the system (e.g. the steady state constraint).

One common objective functions in FBA is maximizing growth or biomass accumulation, which is accomplished by finding the highest possible flux through a biomass equation whose reactants are all the required precursors for making the product: one gram of biomass. Another is minimizing the sum of all fluxes while satisfying a constraint on the rate of biomass accumulation. When maximizing biomass accumulation, if substrate uptake rates (external fluxes) are unbounded, the optimal growth rate is infinite. As mentioned earlier, we apply constraints such that the upper bounds on reactions are not infinite. This makes the maximization of biomass accumulation actually the maximization of biomass given a certain amount of substrate, which is really maximization of efficiency with regards to substrate utilization. Minimization of total flux while satisfying a constraint on biomass accumulation is, likewise, representing a different kind of efficiency. Notably, it has been shown experimentally that the growth rate of *E. coli* evolved under conditions that select for efficient growth becomes very similar to the FBA predicted growth rate [2].

**Some questions you can ask and answer using FBA**

1. You can ask under a given set of growth conditions what genes must carry a non-zero flux in order for the organism to grow (i.e. have positive nonzero biomass flux). This is referred to as “gene essentiality prediction.”
   a. These predictions can be and have been validated experimentally in microbial systems. Typically, these predictions agree with the experimental findings for most genes.
2. You can ask what the maximum yield is for a system you’re using to produce a specific metabolite or set of metabolites and compare this against your actual yield to see if there is room for improvement or if you’re near the theoretical maximum. Moreover, if you are far away from the theoretical maximum, you can probe what the flux map(s) look like that maximize product biosynthesis.
3. You can use FBA as an accounting tool when trying to make an organism produce a new compound. For example, after adding in the heterologous genes necessary to produce a new secondary metabolite, you can use FBA and related techniques to see what fluxes would need to be high to support flux into that metabolite. This may require substantial changes in the overall flux map from that required for regular wild-type biomass accumulation.

**Extreme pathways**

Another way of looking at and exploring the flux solution space is with “extreme pathways.” Imagine we have a three-dimensional space. In order to define any point in that space, we need three coordinates. Similarly, in a higher-dimensional space representing all the flux distributions feasible for a metabolic network, we can define sets of vectors that, in combination with one another, can be used to pinpoint any flux solution. We refer to these individual vectors in these sets as “extreme pathways” because they define the boundaries of the feasible solution space. An appealing aspect of this viewpoint is that these vectors correspond to sets of reactions in the network that are behaving as a unit.

**13C-MFA**

**Reference:** [3]

As highlighted earlier, 13C-MFA is another way of identifying a particular flux map from a possible space of flux maps given the assumption of steady-state. In this case, instead of proposing and optimizing for an objective function, as we did in FBA, we use experimental data to further constrain the solution space, aiming at a single best estimate of fluxes. The data takes the form of steady state labeling measurements.

Recall that we can write mass balance equations for each of the metabolites in our network. Now, without belaboring the theory or the math, we can extend this idea to describe not just the concentrations of each metabolite in our network, but the labeling of each atom in each of these metabolites. In the example network shown in **Figure B5** in [3] we see an example of labeling the C-1 position of glucose. As this labeled glucose makes its way through the fructose-1,6-bisphosphatase reaction, it is split into dihydroxyacetonephosphate (DHAP) and glyceraldehyde-3-phosphate (GAP), with the labeled carbon ending up initially in the C-3 position of DHAP. The ^13^C in DHAP can then move to the C-3 in GAP via the triose phosphate isomerase reaction. ^13^C from both of these triose phosphates can also go back via the reverse fructose-1,6-bisphosphatase reaction, resulting in C-6 labeled hexose molecules (including C-1/C-6 doubly labeled glucose) that are labeled differently than chemically identical ones made directly from the original labeled substrate.

In other words, label tracks through a system and if different pathways or routes through the network feature different rearrangements of atoms which can be labeled with your experimental setup, the distribution of label in the system at isotopic steady state will then reflect the distribution of fluxes.

More specifically, the steady state labeling will tell us the ratios of fluxes through alternative routes in the network which, coupled with constrained (i.e. measured) input and/or output fluxes, will allow us to estimate full flux maps. Please refer to **Box 3. Steady state labeling** in [3] for an example of how to derive the aforementioned ratios from the example network in **Figure B5**.

**Steady State 13C-MFA Examples**

*Developing Brassica napus seeds*

**Referenced paper:** [4,5]

What you need in all flux analysis is an idea of what the metabolic network is. In central metabolism, we can leverage the extensive preexisting biochemical literature to assemble our network model, including enzymes and transporters. This includes information on atom transitions. Some notes:

1. Some chemical species, like CO2, will be produced and consumed by multiple reactions in the metabolic network. Because of this, you get a limited amount of information from some metabolites due to their involvement in multiple processes.
2. On a related note, in eukaryotic systems, metabolites or even entire subnetworks of metabolism can be duplicated across multiple compartments. You will typically not be able to parse out the metabolite pools in these separate compartments, so the signal you get from these compounds will reflect multiple processes.
3. To counteract these difficulties, you should take readouts from as many places as you can. This means, in a system like the developing seed looked at in [4], measuring labeling in sucrose, starch, triacylglycerols, and free and protein-bound amino acids.
4. In [4], the substrates used and their concentrations were set up to mimic the supply of sugars and amino acids supplied to the developing seed by the maternal plant tissue. Additionally, substantial work was done to ensure that the cultured developing embryos resembled normally growing embryos in terms of their growth rate, amino acid and fatty acid profiles, ratios of oil:starch:protein, etc. This was done to ensure that the system was physiologically relevant, which, as we pointed out before, is big challenge (and frequently a very time-consuming one) in flux analysis.

Developing seeds accumulate oil in the form of triacylglycerol (TAG) and this TAG is made mainly of fatty acids built from acetyl-CoA, which itself is derived from hexose phosphates. From six molecules of fructose-6-phosphate, or 36 carbon atoms, we get 12 acetyl-CoA molecules (24 carbon atoms) and 12 CO2 molecules (12 carbon atoms). This suggests that the maximum efficiency with which developing seeds can convert hexose phosphates into TAG is roughly 2/3 of the input carbon. However, if you measure the CO2 emission from developing *B. napus* seeds and compare this to the amount of oil, protein, and cellulose + starch they make, these seeds make oil more efficiently than the apparent 2/3 theoretical maximum.

In [4], it’s shown that a novel route through the metabolic network, which involves recapture of CO2 using RuBisCO without the Calvin-Benson cycle, allows for this higher efficiency. Note that this emphasizes the importance of network thinking. This is not a new pathway *per se*, since all of the constituent reactions were already known as parts of other canonical pathways. Rather, what the flux analysis reveals is a new route passing through parts of glycolysis, the pentose phosphate pathway, and RuBisCO that allows the developing seed to do something novel metabolically.

In contrast to this high efficiency we see in *B. napus*, the closely related crop species *Camelina sativa* has extremely low carbon-conversion efficiency. A similar flux mapping study of developing *C. sativa* embryos was done which showed the reason for this: extremely high fluxes through the oxidative pentose phosphate pathway (OPPP) [5]. This study also showed that the degree to which the OPPP is used and ends up releasing large amounts of CO2 is a function of the light level the seeds are grown in. This highlights the fact that flux maps, rather than representing a concrete and unchanging picture of what the metabolic activity of a given organism or tissue **is**, represent a metabolic phenotype, which is a function of the various environmental and developmental conditions of the organism or organ or cell under consideration.

**References**

1	Y. E. Cheah, J. D. Young Isotopically nonstationary metabolic flux analysis (INST-MFA): putting theory into practice. (2018) *Curr. Opin. Biotechnol.* **54**, 80–87.

2	R. U. Ibarra, J. S. Edwards, B. O. Palsson Escherichia coli K-12 undergoes adaptive evolution to achieve in silico predicted optimal growth. (2002) *Nature*. **420**, 186–189.

3	R. G. Ratcliffe, Y. Shachar-Hill Measuring multiple fluxes through plant metabolic networks. (2006) *Plant J.* **45**, 490–511.

4	J. Schwender, F. Goffman, J. B. Ohlrogge, Y. Shachar-Hill Rubisco without the Calvin cycle improves the carbon efficiency of developing green seeds. (2004) *Nature*. **432**, 779–782.

5	L. M. Carey, T. J. Clark, R. R. Deshpande, J. C. Cocuron, E. K. Rustad, Y. Shachar-Hill High flux through the oxidative pentose phosphate pathway lowers efficiency in developing camelina seeds. (2020) *Plant Physiol.* **182**, 493–506.
