## Supplemental Lesson Plan Day 1 for "Integrative Teaching of Metabolic Modeling and Flux Analysis with Interactive Python Modules"

**Day 01 – Lesson Plan**

**Intended Audience:** Graduate students, postdoctoral researchers, or individuals with equivalent scientific background, in biology, biochemistry, biotechnology or related fields.

**Assumed Prior Knowledge:** Basic familiarity with metabolic processes and biochemistry equivalent to undergraduate level introductory Biochemistry. The lecture notes include linear algebraic and first order differential equations (first year undergraduate or advanced high-school level) but no mathematical manipulations or solutions are involved in the exercises. No knowledge of Python is assumed.

**Resources:**

1. Learners supply their own laptops for carrying out the computational exercises.
2. Pre-installed set of Python packages and libraries, with setup instructions and installation validation for a (non-exclusive) list of hardware and operating system combinations is provided prior to the lesson via a GitHub page: https://github.com/Gibberella/Metabolic-Modeling-Lessons.
3. Jupyter Notebook containing exercises (*Day01_IntroductionToJupyterAndBasicModeling.ipynb*)

**Learning Objectives:**

1. Learn how to use the Python interface that we will be using for the rest of the course and understand the basic outline of how the kinetic simulations are being performed.
2. Analyze perfect and noisy data and use them to relate the values of kinetic parameters in a model of a metabolic network to metabolic simulation results.
3. Analyze the quality of parameter predictions when given more or less informative data points, and relate the information contained in those data to the assumed structure of the metabolic network.
4. Analyze how isotopic labeling data can be used to gain additional information about the operation of a system at steady state.
5. Understand the distinction between isotopic and metabolic steady state.

**Overarching Concepts:**

Key concepts emphasized throughout this workshop include:

- **Concept 1:** The relationship between experimental noise and the confidence one can have in parameter estimates and assumed model architectures.
- **Concept 2:** The distribution of control over fluxes and metabolite concentrations in a network across the reactions in that network.
- **Concept 3:** The uniqueness and identifiability of flux estimates in Flux Balance Analysis and 13C-Metabolic Flux Analysis and their relationship with model complexity.

When these concepts relate to an exercise, we’ll indicate this.

**Content & Exercises**

1.0. (Text section) “***Welcome to Jupyter!”:*** Learners read through an introduction to what Jupyter + Python are and the basic format of the first and subsequent sets of exercises is explained. Learning objectives associated with each set of exercises are also explicitly provided to the learners.

**1.1.**	(Text section + Exercise) **“Using Jupyterlab”:** Learners read through instructions on how to use the Jupyterlab interface. These instructions are interspersed with brief hands-on activities where they print text and use an interactive Python widget to change the output of a function, which they will be doing repeatedly throughout the course.

**1.2.**	(Text section + Exercise) **“Hiding cells”**: Learners are shown how to hide and reveal cells in Jupyterlab.

**2.0.**	(Text section) **“Modeling a metabolic system with first order kinetics”:** Basic theory behind first order kinetic modeling is summarized for learners as review of lecture materials.

**2.1.**	(Text section + Discussion) **“Simulation Logic”:** The text provides a basic outline of the steps our simulation code takes as it simulates the dynamics of a metabolic reaction sequence.

**Discussion Guide:** If the instructor wants course participants to understand the basics of how time-course simulations are run, the code in this section has been extensively annotated to allow a step-by-step rundown of what is happening.

**2.2.**	(Exercise) **“Worked Example”:** Learners are instructed to run some code which then brings up an interactive widget for the first order kinetic simulation. Learners explore modifying the kinetic parameters and note the responses of the concentration, flux, and fractional labeling plots. As they do so, they are asked to consider the following questions:

**(a)** Do the results make sense to you given your parameters?

**(b)** Are there any combinations of parameters whose resulting plots confuse you?

**(c)** Think about the assumptions that went into the model we just constructed. What’s missing? What additional levels of complexity could be added in representing a real biological system.

Learners are encouraged to discuss these questions within their group and to engage the instructor(s) as necessary to understand their results.

**3.0.**	(Text section + Exercise) **“Inferring kinetic parameters from experimental results”:** The text explains to learners that in the real world, we are often interested in inferring parameters from experimental measurements, requiring us to work backwards from our data to infer model architectures and parameters that could generate them. For the exercise, learners are instructed to vary their simulation parameters to come up with a unique set of results and then export these results to a .csv file. Groups then exchange these files with each other and are tasked with fitting their model output to these data by varying the model parameters. Finally, they are asked to share their fitting results with the group with which they swapped data and to consider whether it’s possible for multiple sets of parameter values for the same model can give similar end results, if they were substantially off in their guesses.

**Note:** This requires that learners have a way of exchanging datasets with one another quickly. There are many ways to accomplish this, including creating a class Google Drive or a Discord channel.

**3.1.**	(Text section + Exercise; **Concept 1**) **“Experimental Limitations”:** Similar to the above exercise, except now the simulated data learners export and analyze has synthetic noise added and they only see a subset of the datapoints when fitting. Same as last time, they discuss their results with the group they shared data with. The purpose of this exercise is to demonstrate how the accuracy of parameter estimates relates to the noisiness and sampling density of the underlying dataset one is trying to fit.

**After this activity, learners are given a 10-15 minute break**

**4.0.**	(Text section + Exercise; **Concept 1**) **“Modeling assumptions dictate what information is contained in each datapoint”:** After reading a text section describing how model structure / metabolic network structure informs the information content of specific datapoints, learners engage in an exercise where they try to fit a dataset with limited datapoints. First, learners try fitting the dataset with concentration data for metabolites in the earlier (“upstream”) part of a linear pathway. Then, they try to do the same with metabolites downstream in that same pathway. In this system, the upstream metabolite data provides no information about the downstream kinetic parameters, meaning learners will only be able to get a relatively accurate full set of parameters when they fit using the downstream metabolite datasets.

At the end of the exercise, learners are invited to relate the information content of these datapoints to the structure of the network model that was assumed.

**5.0.**	(Text section + Exercise) **“Steady state kinetics”:** In the text section, learners are introduced to a new model and a review is given of steady state kinetics and the value of isotopic labeling. Learners are asked to consider the limitations of just looking at concentration data. Then, they see a real example of this in the exercise, where they are asked to find multiple sets of parameters that fit a provided dataset of steady-state concentrations. Then, they are given access to the full dataset and can see that their initial guesses didn’t do a good job. Finally, they are given access to the labeling data as well and are asked to consider what additional information it is conveying.

**5.1.**	(Exercise) **“’Steady State’”** In this section, learners are shown a simulation where isotopic and metabolic steady state are reached at different points in the simulation, highlighting the fact that these two states can be decoupled and that this has ramifications for flux analysis.

**Note:** This point is very important for discussions of non-steady-state MFA.
