## Supplemental Lesson Plan Day 2 for "Integrative Teaching of Metabolic Modeling and Flux Analysis with Interactive Python Modules"

**Day 02 – Lesson Plan**

**Intended Audience:** Graduate students / postdoctoral researchers, or individuals with equivalent scientific background, in biology and/or biochemistry.

**Assumed Prior Knowledge:** Basic familiarity with metabolic processes and biochemistry. No knowledge of Python is assumed.

**Resources:**

1. Learners supply their own laptops for carrying out the computational exercises.
2. Pre-installed set of Python packages and libraries, with setup instructions and installation validation provided prior to the lesson via a GitHub page: https://github.com/Gibberella/Metabolic-Modeling-Lessons.
3. Jupyter Notebook containing exercises (*Day02_MetabolicControlAnalysis.ipynb*)

**Learning Objectives:**

1. Analyze the differences between reversible and irreversible first order kinetic models.
2. Observe the increased difficulty/uncertainty involved in fitting to a dataset when additional parameters are added to a model.
3. Carry out metabolic control analysis on a metabolic network and relate flux control coefficients to model parameters.
4. Carry out metabolic control analysis to arrive at estimates of concentration control coefficients.
5. Calculate elasticity values and relate them to the flux and concentration control coefficients calculated in earlier exercises.
6. Observe that the flux control coefficients of all enzymes in a metabolic network add up to a value of 1.

**Overarching Concepts:**

Key concepts emphasized throughout this workshop series include:

- **Concept 1:** The relationship between experimental noise and the confidence one can have in parameter estimates and assumed model architectures.
- **Concept 2:** The uniqueness and identifiability of flux estimates in FBA and 13C-MFA and their relationship with model complexity.
- **Concept 3:** The distribution of control over fluxes and concentrations in a network across the reactions in that network.

When these concepts relate to an exercise, we’ll indicate this.

**Content & Exercises**

1.0. (Text section + Exercise) “**Reversible First Order Kinetics*”:*** Learners read through an introduction to a new model with reversible first order kinetics. They then go through an exercise exploring the effects of varying model parameters. Learners are prompted with: “See how this simulation is similar and different from the ones we ran before. What effect does the inclusion of reverse reactions have?” They are also asked to discuss within their groups: “Do you think this might make fitting parameters harder? Is it possible for our simulation results with reversible reactions to show behavior that you wouldn’t be able to recreate with irreversible first-order kinetics?”

**2.0.**	(Text section + Exercise; **Concept 1**) **“Parameter fitting with reversibility”:** Learners read through a description of how the difficulty of fitting model parameters to data increases the more parameters you have. Learners are then asked to fit a provided dataset. First, they fit only to the concentration data. Then, they fit to the concentration + labeling data. Finally, they can fit to the concentration + labeling + flux values. Learners are asked to consider what information they gained from incorporation of the labeling data, which in this case is the fact that there is a substantial exchange between two metabolite pools that isn’t evident when just analyzing the concentration data.

**3.0.**	(Text section**; Concept 3**) **“Metabolic Control Analysis”**: Learners are given a description of the limitations of thinking about metabolic networks in terms of “rate-limiting” steps. This should be review of the associated lecture material.

**3.1.**	(Text section; **Concept 3**) **“Control over flux through a pathway is distributed across the enzymes in that pathway”:** Learners are provided an overview of flux control coefficient calculations. This is once again review from the lecture material.

**3.2.**	(Exercise + Discussion; **Concept 3**) **“Hands-on MCA Exercise”:** Learners are provided an Excel spreadsheet that designates values for a set of parameters to use for the flux coefficient estimation. They then vary enzyme concentrations up and down in order to estimate control coefficients. After this, the entire class has a discussion where we go through the different groups’ parameters and the control coefficients they arrived at. This is used to spark a conversation about how the different modeled situations affect how control over pathway flux is distributed through the network.

**Note:** The Excel spreadsheet learners need is provided through the GitHub repository.

**Discussion guide:** The whole-class discussion can be facilitated many ways, but we recommend having each group record both the parameter values they changed and their deduced control coefficients on a board visible to all. This allows everyone to relate the parameter changes that were made to the coefficients they calculated. The discussion can be guided to highlight:

1. The distribution of control across multiple enzymes in all cases, which is in sharp contrast to thinking about metabolic pathways and networks in terms of “rate-limiting” or “bottleneck” steps.
2. The relationship between enzyme abundance / activity and control, with reference to a diagram showing the hyperbolic relationship between the activity of an individual enzyme and pathway flux and to the equations describing control coefficients.
3. How increased reversibility of the reaction catalyzed by an enzyme can reduce its control coefficient, but that an enzyme catalyzing a highly reversible reaction can still exert a high degree of control if it is in low abundance.

**After this activity, learners are given a 10-15 minute break**

**3.3.**	(Text Section + Exercise + Discussion**; Concept 3**) **“Concentration Control Coefficients”:** A text section provides a short review of the underlying concept of a concentration control coefficient. Learners use the same parameter values they did in the previous exercise. They then vary enzyme concentrations up and down in order to estimate concentration control coefficients for a specific metabolite. After this, the entire class discusses the different groups’ parameters and the control coefficients they arrived at. We use this to spark a conversation about how different states of the same metabolic network affect how control over the concentration of a specific metabolite is distributed through the network.

**Note:** Similar to **3.2**, having groups record their results on a whiteboard is an effective strategy at this stage.

**Discussion Guide:** Some questions an instructor may consider exploring with learners during the discussion:

1. How are the concentration control coefficients similar and different from the flux control coefficients calculated in **3.2**?
2. When might we care about concentration control coefficients? In what context(s) might these values be useful?
3. If we want to maximize the steady-state concentration of a given metabolite, what kinds of changes to a system might facilitate this based on the exercise results?

**3.4.**	(Text section + Exercise; **Concept 3**) **“Elasticity”:** A text section provides a quick review of the underlying concept of an elasticity. For the exercise, learners are first instructed to use one of the earlier simulations to get estimates of the steady-state concentrations of the various metabolites in our network. They then use these as their starting points to calculate the sensitivity of each flux in the network to the relevant substrate and product metabolites. They do this for each enzyme. Each group has a within-group discussion, and then the whole group has a larger discussion.

**4.0.**	(Text section + Exercise; **Concept 1**) **“Automated MCA”:** Having done a lot of MCA “by-hand”, learners are given an opportunity to get very precise estimates of their concentration control coefficients using an automated procedure, and then told to compare these results against the ones they arrived at earlier to see how close they got.

**Note:** If learners have been shown examples of empirical MCA from the literature, they may have noticed – or the instructor may have highlighted – that these typically involve a very small number of measurements with very different enzyme activities, resulting in a much coarser and less precise measurement of control coefficients than is possible *in silico*.
