## Supplemental Lesson Plan Day 3 for "Integrative Teaching of Metabolic Modeling and Flux Analysis with Interactive Python Modules"

**Day 03 – Lesson Plan**

**Intended Audience:** Graduate students / postdoctoral researchers, or individuals with equivalent scientific background, in biology and/or biochemistry.

**Assumed Prior Knowledge:** Basic familiarity with metabolic processes and biochemistry. No knowledge of Python is assumed.

**Resources:**

1. Learners supply their own laptops for carrying out the computational exercises.
2. Pre-installed set of Python packages and libraries, with setup instructions and installation validation provided prior to the lesson via a GitHub page: <https://github.com/Gibberella/Metabolic-Modeling-Lessons>.
3. Jupyter Notebook containing exercises (*Day03_MichaelisMenten_Updated3rdTime.ipynb*)

**Learning Objectives:**

1. Analyze MCA results in situations where there are variables unaccounted for in the model.
2. Observe how the addition of Michaelis-Menten kinetics to metabolic simulations changes simulation results.
3. Observe the challenges that come with increased model complexity in the presence of experimental noise.
4. Observe how the addition of reversibility to a kinetic simulation with Michaelis-Menten kinetics changes simulation results.
5. Calculate response coefficients and use them to understand the effect of a noncompetitive inhibitor on the flux through a metabolic pathway.
6. Come up with ideas for how metabolic modeling might be incorporated into one’s own research and discuss these with the group.

**Overarching Concepts:**

Key concepts emphasized throughout this workshop series include:

- **Concept 1:** The relationship between experimental noise and the confidence one can have in parameter estimates and assumed model architectures.
- **Concept 2:** The uniqueness and identifiability of flux estimates in FBA and 13C-MFA and their relationship with model complexity.
- **Concept 3:** The distribution of control over fluxes and concentrations in a network across the reactions in that network.

When these concepts relate to an exercise, we’ll indicate this.

**Content & Exercises**

1.0. (Text section; **Concept 3**) “**Metabolic Control Analysis in the Context of Large Metabolic Networks*”:*** Learners read through a description of how metabolic pathways don’t exist in isolation but are embedded in larger metabolic networks. They are then reintroduced to the linear metabolic pathway from the Day 02 MCA exercises.

**1.1.**	(Exercise + Discussion; **Concept 3**) **“Worked Example”:** Learners are asked to once again determine flux control coefficients, *with the difference being that this time there’s actually a branch in the pathway model they are unaware of diverting flux away from product formation*.

**Discussion Guide:** This is a good opportunity to discuss the idea of negative control coefficients and reactions that divert flux away from a desire product.

**1.2.**	(Text section + Exercise + Discussion; **Concept 3**) **“Something Else is Going On”:** The true network architecture is revealed and learners repeat the exercise, this time also calculating the control coefficient for the previously missing enzyme, and then there is another discussion.

**Discussion Guide:** This can naturally lead to a discussion of how data on control coefficients might be used in a biotechnological or chemical engineering context. What do positive and negative control coefficients and their magnitudes tell us about what modifications to a system might be beneficial in achieving a desired goal?

**2.0.**	(Text section) **“Michaelis-Menten Kinetics”:** Learners read through a review of Michaelis-Menten kinetics.

**2.1**	(Text section) **“Adding Michaelis-Menten kinetics to our simulations”:** Learners read details about the implementation of Michaelis-Menten kinetics in the kinetic models.

**2.2.**	(Exercise) **“Simulating with Michaelis-Menten Kinetics”:** Learners are asked to explore different sets of parameter values in a Michaelis-Menten kinetics simulation. They are prompted to find parameter sets that generate results that would be unlikely/impossible to obtain/reproduce using first-order kinetics.

**3.0.**	(Text section + Exercise + Discussion**; Concept 1**) **“Fitting a Complex Reality with a Simple Model”**: Learners are introduced to the idea that there may be cases where they cannot justify a Michaelis-Menten model due to a first-order model approximating its results effectively. Learners then perform an exercise where they attempt to fit data generated by a Michaelis-Menten kinetic simulation using a first order model. They first do this with “perfect” data – that is, no noise added and finely spaced time points. Then they see how their fit looks when noise is added and the data is realistically sparse. After learners have discussed within their groups, they join a whole class discussion on this topic.

**Discussion Guide:** This is a good opportunity to discuss how we must bring a description of the system under consideration to bear in order to model systems and how the complexity of the models we choose are limited by the quality and quantity of data that we have.

**After this activity, learners are given a 10-15 minute break**

**4.0.**	(Text section + Exercise) **“Reversible Michaelis-Menten Kinetics”:** Learners are given a review of reversible Michaelis-Menten kinetics and are then asked to explore the resulting simulated data when using different sets of parameters in a model whose reactions have reversible M-M kinetics.

**5.0.**	(Text section + Exercise + Discussion; **Concept 3**) **“Response Coefficients”:** Student are given an overview of the theory behind response coefficients and introduced to a model where one of the enzymes is subject to noncompetitive inhibition. The learners then carry out an exercise where they explore how the response coefficient is affected by changes in other parameters in the system. They then have an in-group and whole class discussion about conditions that lead to high/low R values and the relationship between the flux control coefficient values for the relevant reaction and response coefficient values.

**6.0**	(Discussion) **“Incorporating metabolic modeling in your work”: ”:** Having extensively explored kinetic modeling concepts and practice, as well as having been exposed to kinetic and constraint-based approaches to metabolic modeling through lecture, learners can consider and then discuss how they might incorporate metabolic modeling into their individual work or to an area of interest to them. These ideas are, ideally, incorporated into the exercises in Day 4, where strengths and limitations of kinetic and constraint-based (Flux Balance Analysis and Metabolic Flux Analysis) are discussed.
