## Supplemental Lesson Plan Day 4 for "Integrative Teaching of Metabolic Modeling and Flux Analysis with Interactive Python Modules"

**Day 04 – Lesson Plan**

**Intended Audience:** Graduate students / postdoctoral researchers, or individuals with equivalent scientific background, in biology and/or biochemistry.

**Assumed Prior Knowledge:** Basic familiarity with metabolic processes and biochemistry. No knowledge of Python is assumed.

**Resources:**

1. Students supply their own laptops for carrying out the computational exercises.
2. Pre-installed set of Python packages and libraries, with setup instructions and installation validation provided prior to the lesson via a GitHub page: https://github.com/Gibberella/Metabolic-Modeling-Lessons.
3. Jupyter Notebook containing exercises (*Day04_ConstraintBasedMethods.ipynb*)

**Learning Objectives:**

1. Analyze a metabolic network using Flux Balance Analysis (FBA), in contrast to kinetic modeling, and understand FBA’s strengths and limitations.
2. Analyze a metabolic network using Flux Variability Analysis (FVA) and random sampling techniques to characterize an FBA solution space.
3. Learn how to work with full genome-scale models as well as reduced core models for FBA analysis.
4. Analyze a metabolic network using MFA, in contrast to kinetic modeling, and understand MFA’s strengths and limitations.
5. Analyze a metabolic network using MFA and an incomplete network specification to understand the ramifications of incorrect network models on flux estimation.
6. Analyze a metabolic network using MFA in the presence of more or less experimental noise to see its impact on the ability to distinguish more or less accurate network models.

**Content & Exercises**

1.0. (Text section) “**Kinetic Modeling in Relation to Constraint-Based Approaches (FBA and MFA)*”:*** Students are reminded of a key limitation of metabolic modeling using kinetic simulations, which is the extensive parameterization necessary to make kinetic simulations work. This should be lecture material review.

**1.1.**	(Text section) **“Steady State Assumption and Constraint-Based Analyses”:** Students are given a review of how constraint-based methods assume metabolic steady-state to make their approach feasible. This is once again a review of lecture material.

**1.2.**	(Exercise + Discussion) **“Kinetic Approach”:** Students use an interactive kinetic simulation widget to obtain the steady state fluxes and fractional labeling of metabolites resulting from a system evolving with time from an arbitrary starting state. Different sets of parameters are used to reach different steady state flux and labeling values. They are asked to record these values for comparison against FBA and MFA results later on.

**Discussion Guide:** In contrast to previous simulations, this simulation records and reports the relative abundances of M0, M1, and M2 mass isomer species of a two-carbon molecule at steady state. Highlighting this and how it relates to the mass spectrometry measurements that are used as input in MFA will be crucial to student understanding.

**2.0.**	(Text section + Exercise + Discussion; **Concept 2**) **“Modeling this system using FBA”:** Students are shown the use of FBA software to simulate the same metabolic network as the one handled in the kinetic exercise of 1.2. Students use different constraints and objective functions to see how these impact their results and whether they make a difference on predictive accuracy. In the end, students have a discussion with the instructors about the strengths and limitations of FBA.

**Discussion Guide:** The concept of degenerate solutions and solution spaces may be unfamiliar to the students, in which case a review of these concepts (introduced in the accompanying lecture material) will be useful to ensure understanding. Asking students to constraint upper and lower bounds for key fluxes to force the solver to generate a “true” flux map that conforms to the kinetic simulation results is a good way of demonstrating that this real solution lies within the solution space, even if it is not the only possible solution given a set of constraints and an objective.

**3.0.**	(Text section; **Concept 2**) **“Flux Variability A and Sampling”:** Brief textual description of the fact that FVA and other sampling approaches can be used to explore solution spaces.

**3.1.**	(Exercise + Discussion; **Concept 2**) **“Flux Variability Analysis”:** Students use the interactive FBA widget to perform FVA and are asked to discuss the results and how to interpret them within their group.

**Discussion Guide:** Flux Variability Analysis is a commonly used technique owing to the lack of unique solutions in FBA. Discuss with the students what kinds of useful information they might get from FVA. For example, how FVA (or a gene-essentiality analysis) can tell you which reactions might feasibly carry zero flux while still allowing an organism to grow.

**3.2.**	(Exercise + Discussion; **Concept 2**) **“Random Sampling”**: Students use the interactive FBA widget to perform random sampling of the solution space and are asked to discuss the results and how to interpret them within their group. Students also see an example of histograms of random sampling results.

**Discussion Guide:** Random sampling provides distributions of possible fluxes for individual reactions as well as random viable whole flux maps. The former can be useful to observe, for example, bimodal or more complex distributions of possible values for individual fluxes, which can be related to qualitative differences between possible flux distributions (e.g. one might observe that there are clusters of solutions featuring high fluxes through the TCA cycle and low fluxes through the OPPP and clusters of solutions featuring high fluxes through the OPPP and low fluxes through the TCA cycle, with each of the reactions in these two pathways showing a bimodal distribution because of this). The latter can be useful if there are metrics or statistics that one wants to calculate from an entire flux map - such as cofactor utilization - and one would like to observe a distribution of those metrics or statistics. These and other points will help students distinguish between random sampling and FVA.

**After this activity, students are given a 10-15 minute break**

**3.3.**	(Exercise + Discussion; **Concept 2**) **“A More Practical FBA Example”**: Students are first shown the features of an Excel spreadsheet of a genome-scale model of *E. coli’s* metabolic network before getting a walkthrough from the instructors on how to read an SBML format model. Then, they generate an FBA prediction of E. coli’s metabolic fluxes with maximal biomass production as the objective function and examine the resulting fluxes. Then, students search the metabolic network file for three specific reactions (citrate synthase, phosphofructokinase, and ATP synthase) and perform FVA for these reactions. From this exercise and explained in the text, it is seen that it is difficult to sample large models, so students switch to using a core model and perform sampling subject to biomass optimization, from which they should observe that

**a.** Given identical inputs, the predicted biomass production rate is different when a core model or genome scale model is used.

**b.** The FVA results from the genome-scale model do not transfer to the core model.

**Discussion Guide:** This is an opportunity to discuss with students the importance of biomass equations to determining FBA flux solutions, as well as the pros and cons of using genome-scale models and core models. In this case, the FVA results differ dramatically due to the availability of alternate routes through the metabolic network in the genome-scale model that simply aren’t available in the core model.

**4.0.**	(Text section + Exercise + Discussion; **Concept 1 & 2**) **“Metabolic Flux Analysis”:** Students are introduced to the necessary input data/model information for the 13C-MFA package *mfapy* and are asked to examine the input files. They then use the steady state labeling values they obtained using the first set of parameters in the earlier kinetic simulation exercise (1.2 above) and put them into the *mfapy* solving procedure to obtain estimated flux values. Once all groups have done this, a whole class discussion is had about the results. The same is done for the 2^nd^ and 3^rd^ sets of parameters.

**Discussion Guide:** Discuss the distinction between net fluxes and forward/reverse fluxes with the students and highlight how in some cases, the forward/reverse fluxes are poorly defined despite accurate net flux estimates. It may also be useful to compare and contrast 13C-MFA with FBA and have a discussion on why 13C-MFA is able to estimate forward and reverse fluxes when FBA is not.

**4.1.**	(Text section + Exercise + Discussion; **Concept 1 & 2**) **“Model Selection”:** Students repeat the kinetic modeling exercise with a new model and record the resulting fluxes and fractional labeling values. Then, they repeat the MFA exercise both with the original, (now incorrect) model and the new, correct model. Students should observe that the model fit is better with the correct model specification.

**Discussion Guide:** Discuss with the students why the model fit looks better when using the correct model versus the incorrect one. Emphasize that incorrect model architectures can lead to label distributions that vary greatly from observations, resulting in higher sum-of-squared-residuals (SSR) for the fit as a whole and/or for particular subsets of labeling values.

**4.2.**	(Exercise + Discussion; **Concept 1 & 2**) **“Model Selection in the Presence of Noise”:** This is a repeat of the above exercise, except students explore applying more or less synthetic noise to their labeling data. They should observe that the addition of even a modest amount of noise results in overlap between the correct and incorrect models in terms of SSR, making model selection more difficult.

**Discussion Guide:** Model selection comes up frequently in 13C-MFA in the context of the compartmentalization of reactions and metabolites in the model network. Providing students with examples of such model selection exercises and how one’s flux map solutions could depend greatly on them would provide valuable insight into the process of doing MFA in the real biological systems. It is also worth emphasizing how, as this exercise demonstrates, your ability to tell a correct from an incorrect model depends on the quality of the input data available.
