## Supplemental Survey Instruments for "Integrative Teaching of Metabolic Modeling and Flux Analysis with Interactive Python Modules"

**Supplemental Material for “Integrative Teaching of Metabolic Modeling and Flux Analysis with Interactive Python Modules”**

Joshua A.M. Kaste, Antwan Green, Yair Shachar-Hill

**Pre-workshop survey**

1. I am a …
   a. Undergraduate
   b. Graduate student
   c. Postdoctoral researcher
   d. Faculty member
   e. None of the above
2. I would describe myself as a …
   a. Biologist
   b. Biochemist
   c. Computational Scientist
   d. None of the above
3. I feel confident in applying and incorporating metabolic modeling techniques to my research question(s)
   a. (Strongly Disagree), (Disagree), (Neutral), (Agree) (Strongly Agree)
4. I feel confident in evaluating the results of a metabolic modeling study or exercise.
   a. (Strongly Disagree), (Disagree), (Neutral), (Agree) (Strongly Agree)
5. I feel confident in identifying metabolic modeling software and techniques that I can apply to my research question(s)
   a. (Strongly Disagree), (Disagree), (Neutral), (Agree) (Strongly Agree)
6. I understand the purpose(s) of metabolic modeling.
   a. (Strongly Disagree), (Disagree), (Neutral), (Agree) (Strongly Agree)
7. I can describe kinetic metabolic modeling and its limitations.
   a. (Strongly Disagree), (Disagree), (Neutral), (Agree) (Strongly Agree)
8. I can describe Metabolic Flux Analysis and its limitations.
   a. (Strongly Disagree), (Disagree), (Neutral), (Agree) (Strongly Agree)
9. I can describe Flux Balance Analysis and its limitations.
   a. (Strongly Disagree), (Disagree), (Neutral), (Agree) (Strongly Agree)
10. I understand the data types I would need to carry out kinetic metabolic modeling.
    a. (Strongly Disagree), (Disagree), (Neutral), (Agree) (Strongly Agree)
11. I understand the data types I would need to carry out Metabolic Flux Analysis.
    a. (Strongly Disagree), (Disagree), (Neutral), (Agree) (Strongly Agree)
12. I understand the data types I would need to carry out Flux Balance Analysis.
    a. (Strongly Disagree), (Disagree), (Neutral), (Agree) (Strongly Agree)
13. I can name the language(s) or software package(s) I would use to incorporate metabolic modeling into my own research
    a. (Strongly Disagree), (Disagree), (Neutral), (Agree) (Strongly Agree)
14. I can critically evaluate the application and results of metabolic modeling in publications and presentations relevant to my area of research.
    a. (Strongly Disagree), (Disagree), (Neutral), (Agree) (Strongly Agree)

**Post-workshop survey**

1. I feel confident in applying and incorporating metabolic modeling techniques to my research question(s)
   a. (Strongly Disagree), (Disagree), (Neutral), (Agree) (Strongly Agree)
2. I feel confident in evaluating the results of a metabolic modeling study or exercise.
   a. (Strongly Disagree), (Disagree), (Neutral), (Agree) (Strongly Agree)
3. I feel confident in identifying metabolic modeling software and techniques that I can apply to my research question(s)
   a. (Strongly Disagree), (Disagree), (Neutral), (Agree) (Strongly Agree)
4. I understand the purpose(s) of metabolic modeling.
   a. (Strongly Disagree), (Disagree), (Neutral), (Agree) (Strongly Agree)
5. I can describe kinetic metabolic modeling and its limitations.
   a. (Strongly Disagree), (Disagree), (Neutral), (Agree) (Strongly Agree)
6. I can describe Metabolic Flux Analysis and its limitations.
   a. (Strongly Disagree), (Disagree), (Neutral), (Agree) (Strongly Agree)
7. I can describe Flux Balance Analysis and its limitations.
   a. (Strongly Disagree), (Disagree), (Neutral), (Agree) (Strongly Agree)
8. I understand the data types I would need to carry out kinetic metabolic modeling.
   a. (Strongly Disagree), (Disagree), (Neutral), (Agree) (Strongly Agree)
9. I understand the data types I would need to carry out Metabolic Flux Analysis.
   a. (Strongly Disagree), (Disagree), (Neutral), (Agree) (Strongly Agree)
10. I understand the data types I would need to carry out Flux Balance Analysis.
    a. (Strongly Disagree), (Disagree), (Neutral), (Agree) (Strongly Agree)
11. I can name the language(s) or software package(s) I would use to incorporate metabolic modeling into my own research
    a. (Strongly Disagree), (Disagree), (Neutral), (Agree) (Strongly Agree)
12. I can critically evaluate the application and results of metabolic modeling in publications and presentations relevant to my area of research.
    a. (Strongly Disagree), (Disagree), (Neutral), (Agree) (Strongly Agree)
13. What did you find useful about the workshop?
    a. Free response
14. What did you not find useful about the workshop?
    a. Free response
15. What changes to the workshop do you think would improve it in future iterations?
    a. Free response

**4-months after survey**

1. If you have had one or more opportunities to apply any of the knowledge you gained in the metabolic modeling workshop, please share your experience(s). If you have not applied any of the knowledge you gained in the metabolic modeling workshop and there are specific reasons why, please share.
   i. Free response
